# σ^28^-dependent small RNA regulation of flagella biosynthesis

**DOI:** 10.1101/2021.08.05.455139

**Authors:** Sahar Melamed, Aixia Zhang, Michal Jarnik, Joshua Mills, Aviezer Silverman, Hongen Zhang, Gisela Storz

## Abstract

Flagella are important for bacterial motility as well as for pathogenesis. Synthesis of these structures is energy intensive and, while extensive transcriptional regulation has been described, little is known about the posttranscriptional regulation. Small RNAs (sRNAs) are widespread posttranscriptional regulators, most base pairing with mRNAs to affect their stability and/or translation. Here we describe four UTR-derived sRNAs (UhpU, MotR, FliX and FlgO) whose expression is controlled by the flagella sigma factor σ^28^ (*fliA*) in *Escherichia coli*. Interestingly, the four sRNAs have varied effects on flagellin protein levels, flagella number and cell motility. UhpU, corresponding to the 3’ UTR of a metabolic gene, likely has hundreds of targets including a transcriptional regulator at the top flagella regulatory cascade connecting metabolism and flagella synthesis. Unlike most sRNAs, MotR and FliX base pair within the coding sequences of target mRNAs and act on ribosomal protein mRNAs connecting ribosome production and flagella synthesis. The study shows how sRNA-mediated regulation can overlay a complex network enabling nuanced control of flagella synthesis.

## Introduction

Most bacteria are motile and can swim through liquid and semiliquid environments in large part driven by the flagellum. The highly complex bacterial flagellum consists of three major domains: an ion-driven motor, which can provide torque in either direction; a universal joint called the hook-basal body, which transmits motor torque; and a 20 nm thick hollow filament tube composed of the flagellin subunit, which acts as a propeller (reviewed in (Altegoer & Bange, 2015, Nakamura & Minamino, 2019)). The complete flagellum is comprised of many proteins, and the flagellar regulon encompasses more than 50 genes. Since it is estimated that one flagellum constitutes ∼2% of the total protein in the cell, synthesis is a costly process requiring extensive use of ribosomes (reviewed in (Soutourina & Bertin, 2003, Guttenplan & Kearns, 2013)).

To ensure that flagellar components are made in the order in which they are needed, transcription of the genes in the regulon is activated in a sequential manner in *Escherichia coli* (Kalir *et al*., 2001) and *Salmonella enterica* (reviewed in (Chevance & Hughes, 2008)). The genes can be divided into three groups based on their time of activation: early genes, middle genes, and late genes (*Figure 1A*). The FlhDC transcription regulators, encoded by the two early genes, activate the transcription of the middle genes (Class 2), which are required for the hook-basal body. FlhDC also activates transcription of *fliA*, encoding sigma factor σ^28^ (Fitzgerald *et al*., 2014). σ^28^ in turn activates transcription of the late genes responsible for completing the flagellum and the chemotaxis system (Class 3). σ^28^ additionally increases expression of several of the middle genes (Class 2/3) (Fitzgerald *et al*., 2014). σ^28^ activity itself is negatively regulated by the anti-sigma factor, FlgM, which is transported out of the cell when the hook-basal body complex is complete (reviewed in (Smith & Hoover, 2009, Osterman *et al*., 2015)). Given the numerous components required at different times and in different stoichiometries during flagellum assembly, various factors can be rate limiting under specific conditions (reviewed in (Chevance & Hughes, 2008)). The dependence of flagella synthesis on FlhDC and σ^28^ generates a coherent feed-forward loop. In this loop, the first regulator (FlhDC) activates the second regulator (σ^28^), and they both additively activate their target genes. This results in prolonged flagellar expression, protecting the flagella synthesis from a transient loss of input signal (Kalir *et al*., 2005).

**Figure 1.**
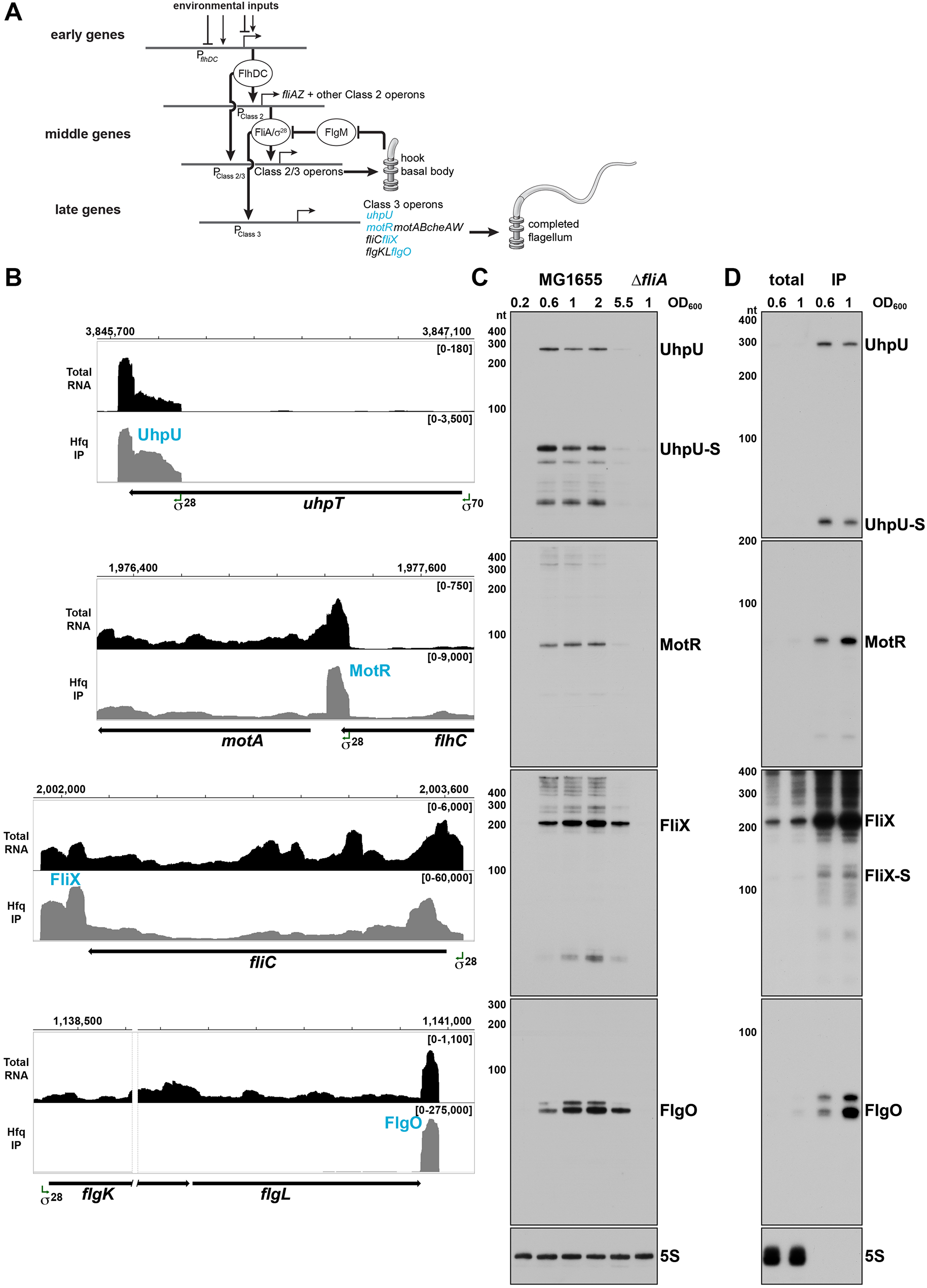
σ^28^-dependent sRNAs are primarily expressed in log phase. (**A**) Overview of the flagellar regulon. The early genes initiate the transcription of the middle genes, including *fliA* which encodes σ^28^. In turn, σ^28^ initiates the transcription of the late genes and enhances the transcription of some of the middle genes. For the middle and late genes, only selected operons are shown. The sRNAs analyzed in this study are colored in blue. This model was inspired by (Kalir *et al*., 2005). (**B**) Browser images showing levels of UhpU, MotR, FliX and FlgO sRNAs in total RNA (black) and Hfq co-immunoprecipitation (gray) libraries. Normalized read count ranges are shown in the upper right of each frame. Data analyzed is from (RIL-seq experiment 1, (Melamed *et al*., 2020). (**C**) Northern blot analysis of total RNA from WT (GSO983) or Δ*fliA* (GSO1068) cells grown to the indicated time points. The same membrane was probed for all four σ^28^-dependent sRNAs. A full-length transcript (∼260 nt) and several processed transcripts, of which one is predominant (UhpU-S, ∼60 nt), are detected for UhpU, one prominent band (∼95 nt) is detected for MotR, one prominent band (∼200 nt) is detected for FliX, and two close bands close in size (∼75 nt) are detected for FlgO. (**D**) WT (GSO983) cells were grown to OD_600_ ∼0.6 and ∼1.0. RNA was extracted from total lysates as well as samples from co-immunoprecipitation with Hfq, separated on an acrylamide gel, transferred to a membrane, and probed for σ^28^-dependent sRNAs. A ∼100 nt FliX band (FliX-S) was revealed immunoprecipitating with Hfq. In (**C**) and (**D**), RNAs were probed sequentially on the same membrane, and the 5S RNA served as a loading control. **Figure supplement 1.** Sequences and predicted structures of UhpU, MotR, FliX and FlgO sRNAs and effect of carbon source on sRNA levels. **Figure supplement 2.** UhpU, MotR, FliX and FlgO levels across growth.

Given flagella are so costly to produce, synthesis is tightly regulated such that flagellar components are only made when motility is beneficial. Thus, flagellar synthesis is strongly impacted by environmental signals. For instance, flagellar gene expression is decreased in the presence of D-glucose, in high temperatures, high salt, and extreme pH, as well as the presence of DNA gyrase inhibitors (Shi *et al*., 1993, Adler & Templeton, 1967). The flagellar genes are activated under oxygen-limited conditions (Landini & Zehnder, 2002) and at various stages of infection (reviewed in (Erhardt, 2016)). Consequently, transcription of many genes in the flagellar regulon is regulated in response to a range of environmental signals. For example, transcription of *flhDC* is controlled by at least 13 transcription factors, each of them active under different conditions (reviewed in (Prüß, 2017)).

While the activation of flagella synthesis has been examined in some detail, there has been less investigation into the termination of synthesis, which we presume is equally important for the conservation of resources. Additionally, while transcriptional regulation of flagella genes has been studied for many years, the post-transcriptional control of the regulon has only received limited attention. Small RNAs (sRNAs) that can originate from many different genetic loci (reviewed in (Adams & Storz, 2020)) are key post-transcriptional regulators in bacteria. They usually regulate their targets in trans via limited base-pairing, affecting translation and/or mRNA stability (reviewed in (Hör *et al*., 2020)). Many characterized sRNAs are stabilized and their base pairing with targets increased by RNA chaperones, of which the hexameric, ring-shaped Hfq protein has been studied most extensively (reviewed in (Updegrove *et al*., 2016, Holmqvist & Vogel, 2018)). The only post-transcriptional control by base pairing sRNAs described for the *E. coli* flagellar regulon thus far is negative regulation of *flhDC* by ArcZ, OmrA, OmrB, OxyS (De Lay & Gottesman, 2012), and AsflhD (encoded antisense to *flhD*)(Lejars *et al*., 2022), positive regulation of the same mRNA by McaS (Thomason *et al*., 2012), and negative regulation of *flgM* by OmrA and OmrB (Romilly *et al*., 2020). These sRNAs and a few other sRNAs also were shown to affect motility and biofilm formation (Bak *et al*., 2015).

In this study, we characterized four σ^28^-dependent sRNAs, which were detected with their targets on Hfq through RIL-seq methodology that captures the sRNA-target interactome (Melamed *et al*., 2016, Melamed *et al*., 2020). These sRNAs originate from the untranslated regions (UTRs) of mRNAs, three of which belong to the flagellar regulon. We identified a wide range of targets for the sRNAs, including genes related to flagella and ribosome synthesis and observed that the sRNAs act on some of these targets by unique modes of action. We also found that three of these sRNAs regulate flagella number and bacterial motility, possibly imposing temporal control on flagella synthesis and integrating metabolic signals into this complex regulatory network.

## Results

### σ^28^-dependent sRNAs are expressed sequentially in log phase cells

Analysis of several different RNA-seq data sets suggested the expression of four σ^28^-dependent sRNAs in *E. coli*. σ^28^-dependent expression of the sRNAs was detected using ChIP-seq and RNA-seq in a comprehensive analysis of the σ^28^ regulon (Fitzgerald *et al*., 2014), while the position and nature of the 5’ ends were revealed by a 5’ end mapping study (Thomason *et al*., 2015). Regulatory roles were indicated by binding to other RNAs in RIL-seq data (Melamed *et al*., 2016, Melamed *et al*., 2020, Bar *et al*., 2021). The four sRNAs originate from the UTRs of protein coding genes (*Figure 1B* and *Figure 1—figure supplement 1A*). UhpU corresponds to the 3’ UTR of *uhpT*, which encodes a hexose phosphate transporter (Marger & Saier, 1993). UhpU is transcribed from its own promoter inside the coding sequence (CDS) of *uhpT* (Thomason *et al*., 2015). The other three σ^28^-dependent sRNAs correspond to the UTRs of the late genes in the flagellar regulon. MotR originates from the 5’ UTR of *motA*, which encodes part of the flagellar motor complex. Based on previous transcription start site analysis, the promoter for *motR* is within the *flhC* CDS and is also the promoter of the downstream *motAB-cheAW* operon (Thomason *et al*., 2015, Fitzgerald *et al*., 2014). FliX originates from the 3’ UTR of *fliC,* which encodes flagellin, the core component of the flagellar filament (reviewed in (Thomson *et al*., 2018)). FlgO originates from the 3’ UTR of *flgL*, a gene that encodes a junction protein shown to connect the flagella to the hook in *S. enterica* (Ikeda *et al*., 1987). The observation that FliX and FlgO levels decline substantially in RNA-seq libraries treated with 5’ phosphate-dependent exonuclease to deplete processed RNAs (Thomason *et al*., 2015), indicates that both of these sRNAs are processed from their parental mRNAs.

Northern blot analysis confirmed σ^28^-dependent synthesis of these sRNAs since expression was significantly decreased in a mutant lacking σ^28^ (Δ*fliA*) (*Figure 1C*). Given that most σ^28^-dependent mRNAs encode flagella components, the regulation suggests the sRNAs impact flagella synthesis. The northern analysis also showed that the levels of the four σ^28^-dependent sRNAs are highest in the transition from mid-exponential to stationary phase growth, though there are some differences with UhpU and MotR peaking before FliX and FlgO (*Figure 1C* and *Figure 1—figure supplement 2*). Since flagellar components are expressed at precise times, the difference in the UhpU and MotR peak times compared to the FliX and FlgO peak times hints at different roles for each of these sRNAs. For UhpU, two predominant bands were observed, a long transcript and a shorter transcript processed from UhpU (denoted UhpU-S), which corresponds to the higher peak in the sequencing data (*Figure 1B*). One prominent band was detected for MotR and for FliX, while a doublet was observed for FlgO. Additional higher bands detected by the MotR probe could be explained by RNA polymerase read through of the MotR terminator into the downstream *motAB-cheAW* operon, while the additional bands seen for FliX could be explained by alternative processing of the *fliC* mRNA.

We also examined the levels of the four sRNAs in minimal media (M63) supplemented with different carbon sources (*Figure 1—figure supplement 1B*). Generally, the sRNAs levels in minimal medium are comparable to or slightly higher to the levels in rich media (LB) except in medium with glucose-6-phosphate (G6P), where the levels of UhpU-S are significantly elevated while the levels of full-length UhpU transcript and the other σ^28^-dependent sRNAs are decreased. These observations suggest an alternative means for UhpU-S generation from the *uhpT* mRNA known to be induced by G6P (Verhamme *et al*., 2001). We also observe more FliX products, particularly for cells grown in minimal medium with ribose or galactose.

The predicted structures for the four σ^28^-dependent sRNAs (*Figure 1—figure supplement 1C*), with strong stem-loops at the 3’ ends, are consistent with the structures of known Hfq-binding sRNAs and the association with Hfq observed in the RIL-seq data (Melamed *et al*., 2016). To confirm Hfq binding, we probed RNA that co-immunoprecipitated with Hfq (*Figure 1D*). Strong enrichment and fewer background bands were observed for all of the sRNAs; ∼260 nt and ∼60 nt bands for UhpU and UhpU-S, respectively, a ∼95 nt band for MotR, a ∼200 nt band for FliX and a doublet of ∼75 nt bands for FlgO. For FliX, we also detected a second ∼100 nt FliX band (denoted FliX-S; *Figure 1—figure supplement 1A*) that corresponds to the 3’ peak in the sequencing data (*Figure 1B*) and includes one of the repetitive extragenic palindromic (REP) sequences downstream of *fliC*.

### σ^28^-dependent sRNAs impact flagella number and bacterial motility

To begin to decipher the roles of the four σ^28^-dependent sRNAs, we constructed plasmids for overexpression of the sRNAs (*Figure 2—figure supplement 1A*). Given that it was challenging to obtain constructs constitutively overexpressing UhpU because all clones had mutations, this sRNA could only be expressed from a plasmid when controlled by an IPTG-inducible P_lac_ promoter (Guo *et al*., 2014), hinting at a critical UhpU role in *E. coli* vitality. The other sRNAs were expressed from a plasmid with the constitutive P_LlacO-1_ promoter (Urban & Vogel, 2007). We also obtained a plasmid constitutively overexpressing MotR*, a more abundant derivative of MotR identified by chance (TGC at positions 6-8 mutated to GAG; *Figure 1—figure supplement 1A*).

We tested the effects of overexpressing the sRNAs on flagellar synthesis by determining the number of flagella by electron microscopy (EM) and on bacterial motility by assaying the spread of cells on 0.3% agar plates. The WT *E. coli* strain used throughout the paper is highly motile due to an IS1 insertion in the *crl* gene (*crl*^-^), thus eliminating expression of a protein that promotes σ^S^ binding to the RNA polymerase core enzyme (Typas *et al*., 2007), and resulting in higher expression of the flagellar regulatory cascade (Pesavento *et al*., 2008). However, we also assayed a less motile strain with the restored *crl*^+^ gene for UhpU and MotR effects on motility, given that no effects were observed with the highly motile *crl*^-^ strain.

Intriguingly, overexpression of the individual sRNAs had different consequences. UhpU overexpression caused a slight increase in flagella number (*Figure 2A*) and a marked increase in motility (*Figure 2B*). Overexpression of MotR, particularly MotR*, led to a dramatic increase in the flagella number (*Figure 2C* and *Figure 2—figure supplement 2A*) and MotR but not MotR* had a slight effect on motility (*Figure 2D* and *Figure 2—figure supplement 2B*). It has been suggested that the run/tumble behavior of bacteria, which affect their swimming, is only weakly dependent on number of flagella (Mears *et al*., 2014), possibly explaining these somewhat contradictory effects on flagella number and motility. In contrast to UhpU and MotR, FliX overexpression led to a reduction in the number of flagella (*Figure 2E*), an effect that was even more pronounced in a strain overexpressing FliX-S (*Figure 2—figure supplement 2C*). Overexpression of FliX-S but not FliX also reduced bacterial motility (*Figure 2F* and *Figure 2— figure supplement 2D*). While FliX-S overexpression seems to lead to aflagellated bacteria, we hypothesize that the sRNA is delaying but not eliminating flagella gene expression, explaining why the bacteria are still moderately motile. Some motility phenotypes can be explained by differences in growth rate, but we do not think that this is the case for MotR and FliX as we observed only slight effects on growth upon MotR, MotR*, FliX and FliX-S overexpression (*Figure 2—figure supplement 1B*). FlgO overexpression did not result in detectable changes in our assays (*Figure 2G* and *Figure 2H*). Together, these results show that the σ^28^-dependent sRNAs have a range of effects on flagella number and motility, with UhpU and MotR, which are expressed first, increasing both phenotypes and FliX, which is expressed later, decreasing both. Given that MotR* and FliX-S have stronger effects for some phenotypes and provide a bigger dynamic range, these derivatives were included in subsequent assays.

**Figure 2.**
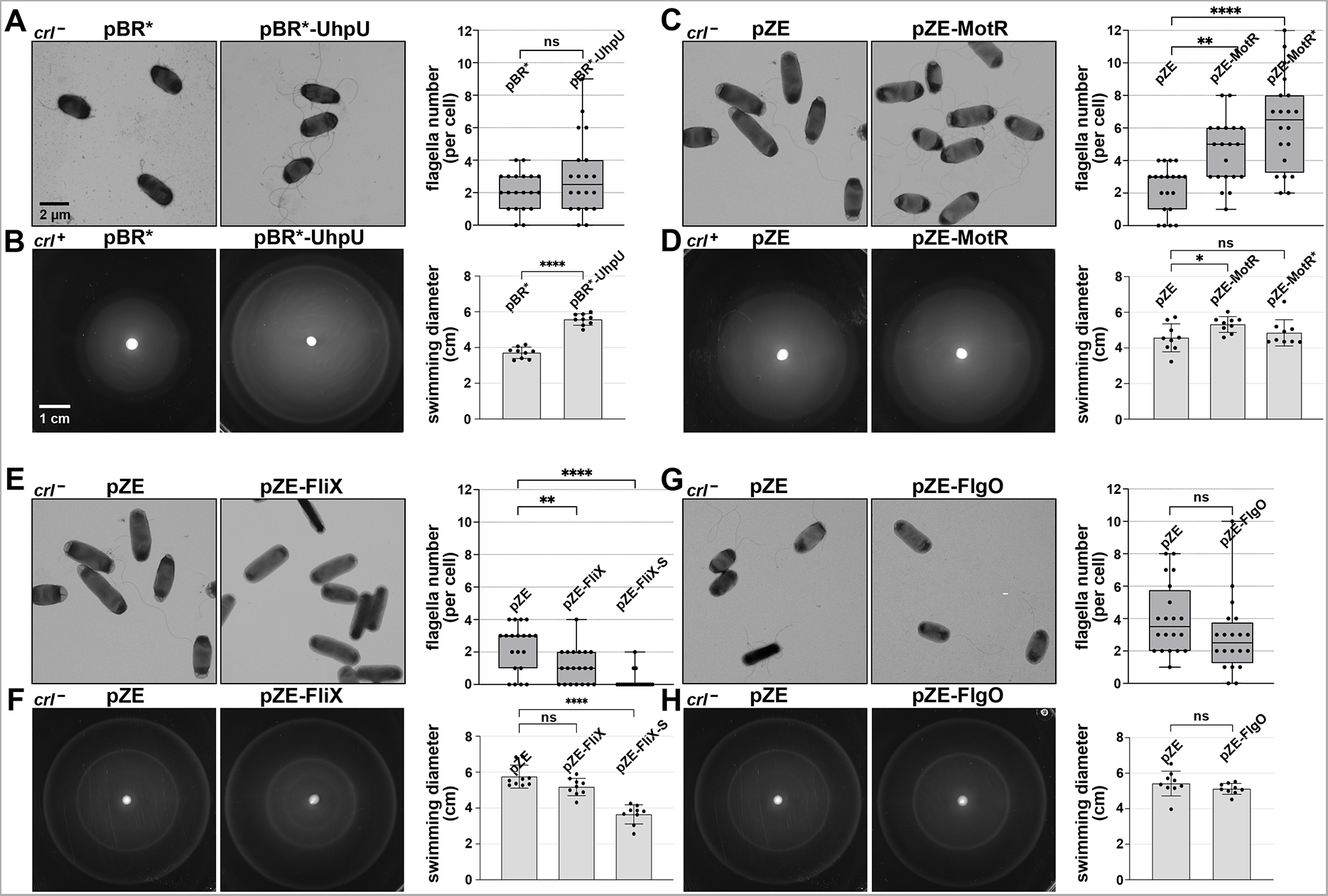
Overexpression of the σ^28^-dependent sRNAs leads to differences in flagella number and motility. (**A**) Moderate increase in flagella number with UhpU overexpression based on electron microscopy analysis for WT (*crl*^-^) cells carrying an empty vector or overexpressing UhpU. (**B**) Increased motility with UhpU overexpression based on motility in 0.3% agar for WT (*crl*^+^) cells carrying an empty vector or overexpressing UhpU. (**C**) Increase in flagella number with MotR overexpression based on electron microscopy analysis for WT (*crl*^-^) cells carrying an empty vector or overexpressing MotR. (**D**) Slight increase in motility with MotR overexpression based on motility in 0.3% agar for WT (*crl*^+^) cells carrying an empty vector or overexpressing MotR. (**E**) Reduction in flagella number with FliX overexpression based on electron microscopy analysis for WT (*crl*^-^) cells carrying an empty vector or overexpressing FliX. (**F**) Reduced motility with FliX overexpression based on motility in 0.3% agar for WT (*crl*^-^) cells carrying an empty vector or overexpressing FliX. (**G**) No change in flagella number with FlgO overexpression based on electron microscopy analysis for WT (*crl*^-^) cells carrying an empty vector or overexpressing FlgO. (**H**) No change in motility with FlgO overexpression based on motility in 0.3% agar for WT (*crl*^-^) cells carrying an empty vector or overexpressing FlgO. Cells in (**A**) and (**B**) were induced with 1 mM IPTG. Quantification for all the assays is shown on the right. For (**A**), (**C**), (**E**) and (**G**) quantification of the number of flagella per cell was done by counting the flagella for 20 cells (black dots), and a one-way ANOVA comparison was performed to calculate the significance of the change in flagella number (ns = not significant, ** = P<0.01, **** = P<0.0001). Each experiment was repeated three times and one representative experiment is shown. The bottom and top of the box are the 25th and 75th percentiles, the line inside the box is the median, the lower and the upper whiskers represent the minimum and the maximum values of the dataset, respectively. While some differences in cells size and width were observed in the EM analysis, they were not statistically significant. The experiments presented in (**C**) and (**E**) were carried out on same day, and the same pZE sample is shown. Graphs for (**B**), (**D**), (**F**) and (**H**) show the average of nine biological repeats. Error bars represent one SD, and a one-way ANOVA comparison was performed to calculate the significance of the change in motility (ns = not significant, * = P<0.05, **** = P<0.0001). The scales given in (**A**) and (**B**) are the same for all EM images and all motility plates, respectively. **Figure supplement 1.** Expression of σ^28^-dependent sRNAs from plasmids and the effect of MotR and FliX overexpression on growth. **Figure supplement 2.** Effects of MotR* and FliX-S overexpression on flagella number and motility.

### σ^28^-dependent sRNAs have wide range of potential targets based on RIL-seq analysis

To understand the phenotypes associated with overexpression of the σ^28^-dependent sRNAs expression, we took advantage of the sRNA-target interactome data obtained by RIL-seq (Melamed *et al*., 2020, Melamed *et al*., 2016). We analyzed the data (*Supplementary file 1*) generated from 18 samples representing six different growth conditions, which included different stages of bacterial growth in rich medium as well as growth in minimal medium and iron limiting conditions. We selected targets for further characterization if they were detected for at least four different conditions. The sRNAs differ significantly in their target sets (*Figure 3—figure supplement 1A*). In general, UhpU is a hub with hundreds of RIL-seq targets. Its target set comprises a wide range of genes, including multiple genes that have roles in flagella synthesis and carbon metabolism. MotR and FliX were associated with fewer targets, but intriguingly, both sets were enriched for genes encoding ribosomal proteins. We also noted that the *fliC* gene encoding flagellin was present in the target sets for UhpU, MotR, and FliX. Although FlgO is one of the most strongly enriched sRNAs upon Hfq purification (ranked fourth in Melamed et al., 2020), it had the smallest set of targets. Almost none of the targets were found in more than two conditions and only *gatC* was detected in four conditions, hinting FlgO might not act as a conventional Hfq-dependent base-pairing sRNA. Unlike for most characterized sRNA targets, the RIL-seq signal for the sRNA interactions with *fliC* and the ribosomal protein genes is internal to the CDSs (*Supplementary file 1* and *Figure 3—figure supplement 1B*). Before turning our attention to these unique targets, we first examined the UhpU interaction with a canonical target.

### UhpU represses expression of the LrhA transcriptional repressor of *flhDC*

We were intrigued to find that the mRNA encoding the transcription factor LrhA, which represses *flhDC* transcription, was among the top RIL-seq interactors for UhpU (*Supplementary file 1*). The signals that activate this LysR-type transcription factor (Lehnen *et al*., 2002), are not known, but the *lrhA* mRNA has an unusually long 371 nt 5’ UTR (*Figure 3A*), a feature that has been found to correlate with post-transcriptional regulation (reviewed in (Adams & Storz, 2020)). The predicted base pairing between UhpU and the *lrhA* 5’-UTR (*Figure 3B*) corresponds to the seed sequence suggested for UhpU (Melamed *et al*., 2016).

**Figure 3.**
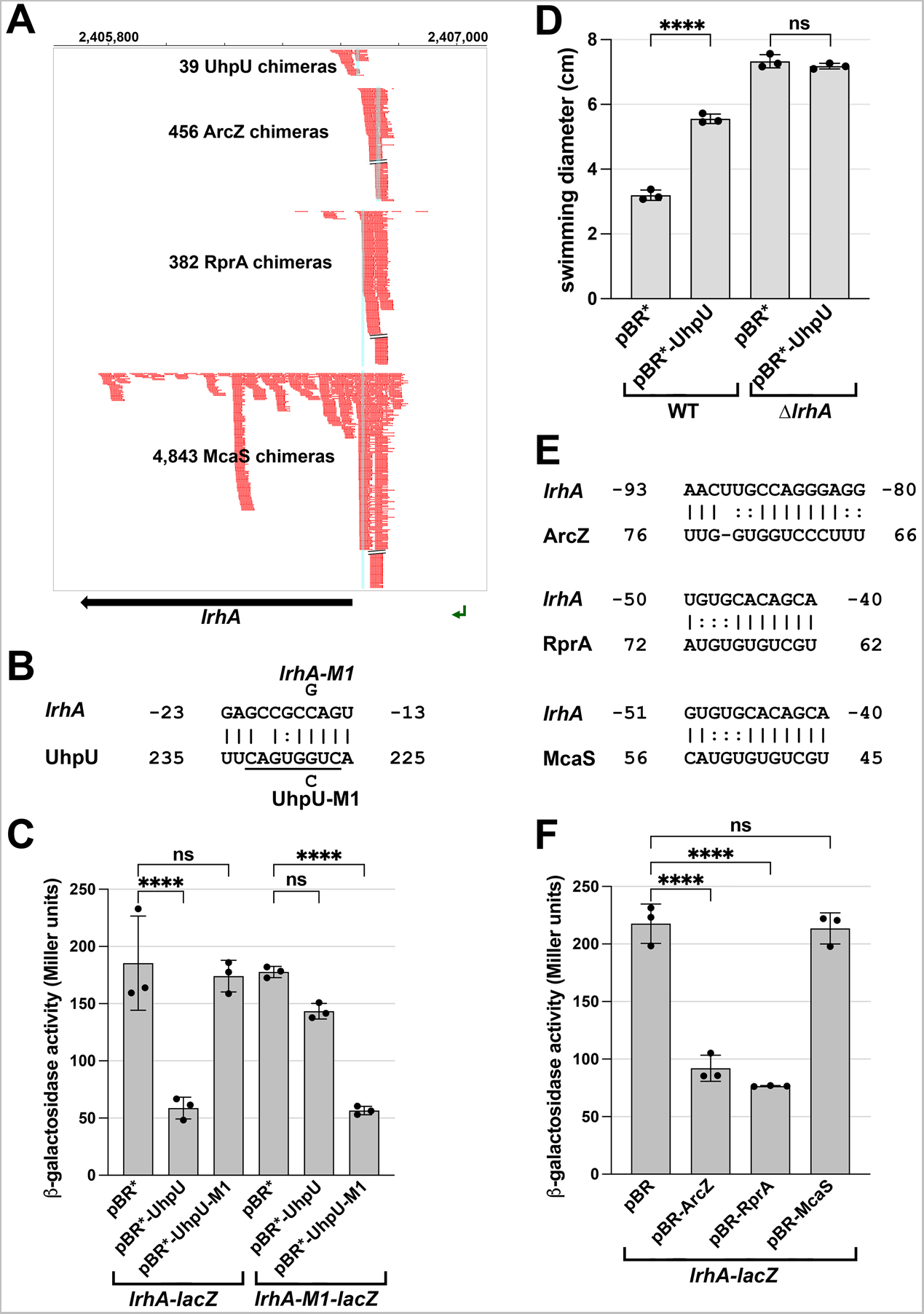
Multiple sRNAs repress LrhA synthesis. (**A**) Browser image showing chimeras (in red) for UhpU, ArcZ, RprA and McaS, at the 5’ UTR region of *lrhA*. Blue highlighting indicates position of sRNA-*lrhA* base pairing. Data analyzed is from (Melamed *et al*., 2020). (**B**) Base-pairing between *lrhA* and UhpU with sequences of mutants assayed. Seed sequence predicted by (Melamed *et al*., 2016) is underlined. Numbering is from AUG of *lrhA* mRNA and +1 of UhpU sRNA. (**C**) UhpU represses *lrhA-lacZ* fusion. β-galactosidase assay detecting the levels of *lrhA-lacZ* and *lrhA-M1-lacZ* translational fusions in response to UhpU and UhpU-M1 overexpression. (**D**) UhpU does not affect motility when LrhA is absent, based on motility in 0.3% agar for WT (*crl^+^*) cells or Δ*lrhA* cells (GSO1179) carrying an empty vector or overexpressing UhpU. Graph shows the average of three biological repeats, and error bars represent one SD. One-way ANOVA comparison was performed to calculate the significance of the change in motility (ns = not significant, **** = P<0.0001). (**E**) Predicted base-pairing between *lrhA* and ArcZ, RprA or McaS. Numbering is from AUG of *lrhA* mRNA and +1 of indicated sRNAs. (**F**) Down regulation of *lrhA* by ArcZ and RprA but not McaS. β-galactosidase assay detecting the levels of *lrhA-lacZ* translational fusions in response to ArcZ, RprA and McaS overexpression. For (**C**) and (**F**), graphs show the average of three biological repeats, and error bars represent one SD. One-way ANOVA comparison was performed to calculate the significance of the change in β-galactosidase activity (ns = not significant, **** = P<0.0001). **Figure supplement 1.** Interactomes for σ^28^-dependent sRNAs.

To test the effects of UhpU on this target, we fused the 5’ UTR of *lrhA*, which includes the region of the RIL-seq *lrhA*-UhpU chimeras and the predicted base-pairing region, to a *lacZ* reporter (Mandin & Gottesman, 2009). UhpU overexpression reduced expression of the chromosomally-encoded P_BAD_-*lrhA*-*lacZ* reporter (*Figure 3C*). A single nucleotide mutation in the base pairing region of *uhpU* (*uhpU-M1*) eliminated UhpU repression of *lrhA-lacZ*, while a complementary mutation introduced into the chromosomal *lrhA*-*lacZ* fusion (*lrhA-M1*) restored the repression providing direct evidence for UhpU base pairing to *lrhA* leading to repression. Down-regulation of LrhA by UhpU, which is expected to lead to increased FlhDC levels, is in accord with the positive impact of UhpU on motility (*Figure 2*). To test this model, we monitored the effect of UhpU on bacterial motility in a *lrhA* deletion strain compared to a WT strain (*Figure 3D*). With UhpU overexpression, motility was increased in the WT background as expected. In contrast, while the Δ*lrhA* strain was more motile, likely due to *flhDC* de-repression, motility was unaltered by high levels of UhpU indicating that significant UhpU effects on motility are mediated by LrhA.

Interestingly, the RIL-seq data also suggested that *lrhA* directly interacts with other sRNAs such as ArcZ, RprA and McaS (*Figure 3A*). Regions of predicted base pairing overlap known seed regions for these sRNAs (*Figure 3E*). In translational reporter assays using the *lrhA-lacZ* fusion, both RprA and ArcZ reduced expression, while McaS, despite having the most chimeras, had no effect (*Figure 3F*). Possibly the McaS-*lrhA* interaction has other regulatory consequences such as McaS inhibition. Intriguingly, ArcZ, RprA and LrhA form a complex regulatory network with the general stress response sigma factor σ^S^ encoded by *rpoS*, as previous studies showed that LrhA represses the expression of *rprA* and *rpoS* (Peterson *et al*., 2006), while ArcZ and RprA increase *rpoS* expression (reviewed in (Mika & Hengge, 2014)).

### UhpU, MotR and FliX modulate flagellin levels

The high number of chimeras between UhpU, MotR or FliX with the *fliC* mRNA encoding flagellin were striking, particularly between the 3’ end of *fliC* corresponding to FliX (blue) and the 5’ end of *fliC* (red) (*Figure 4A*). As mentioned above, it was also noteworthy that most of the chimeras were internal to the *fliC* CDS. When we examined the consequences of overexpressing UhpU, MotR, MotR*, FliX or FliX-S on the levels of the flagellin protein, we observed somewhat increased levels of flagellin, both as cytosolic monomers (*Figure 4B*) and de-polymerized flagella (*Figure 4—figure supplement 1A*) with UhpU and MotR* overexpression and reduced levels with FliX or FliX-S overexpression. These differences are reflected in increased levels of the *fliC* mRNA with overexpression of UhpU, particularly in a *crl*^+^ background, or MotR or MotR*, particularly at OD_600_ ∼0.2 (*Figure 4C* and *Figure 4—figure supplement 1B*). In contrast, *fliC* mRNA levels decreased with FliX and FliX-S overexpression (*Figure 4C* and *Figure 4—figure supplement 1B*). In general, the impacts of the sRNAs on flagellin protein and *fliC* mRNA levels are consistent with the increased flagella number and/or motility upon UhpU or MotR overexpression and decreased flagella number upon FliX overexpression. Comparatively, the effects of MotR and MotR* on flagella number and *fliC* mRNA levels were stronger than the effects on the flagellin protein; possibly increases in flagellin levels are masked by the abundance of the protein.

**Figure 4.**
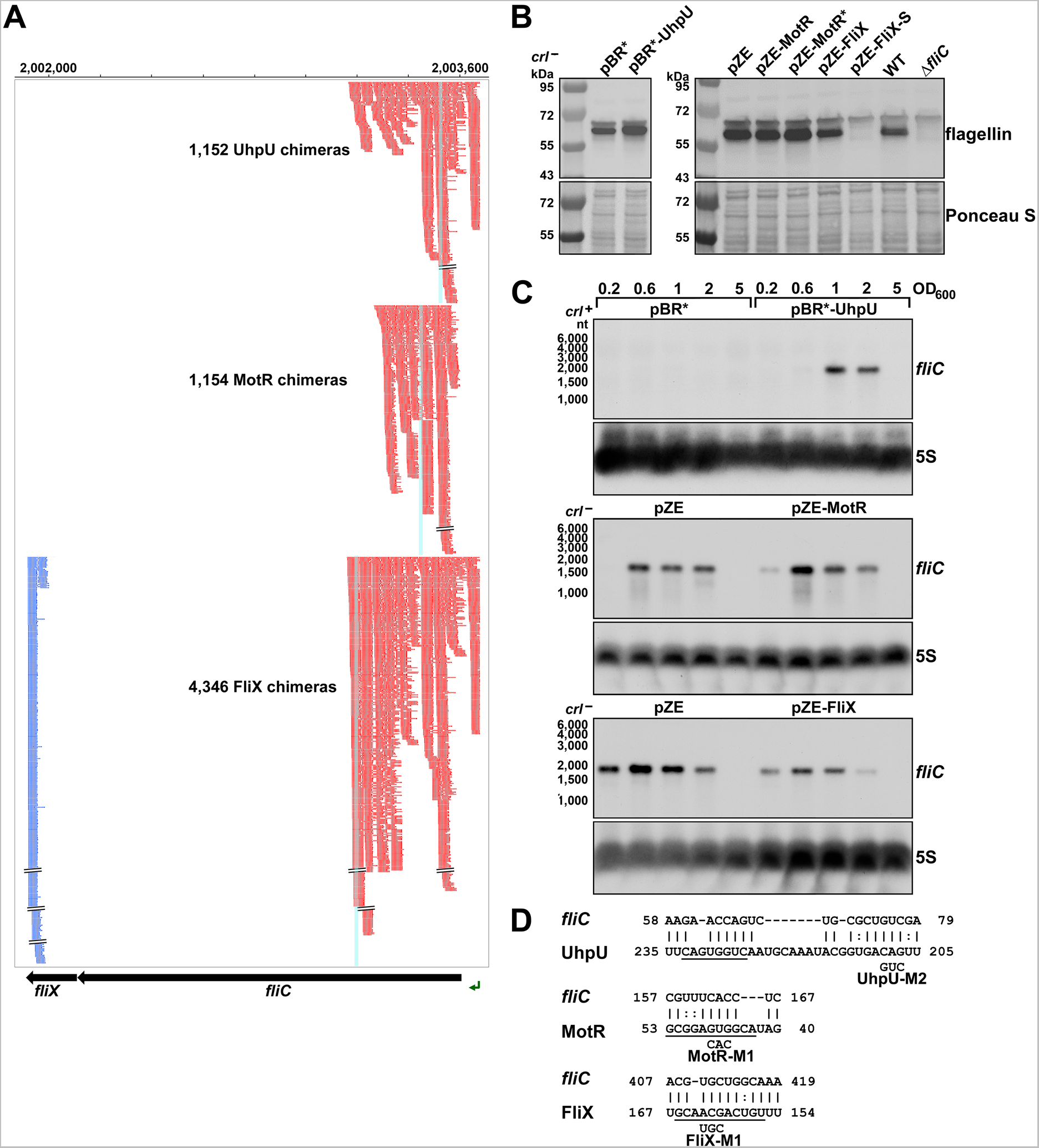
Multiple sRNAs regulate flagellin synthesis. (**A**) Browser image showing chimeras (red and blue) for UhpU, MotR, and FliX at the *fliCX* region. Data analyzed is from (RIL-seq experiment 1, (Melamed *et al*., 2020). Red and blue lines indicate the RNA in the region is first or second RNA in the chimera, respectively. Blue highlighting indicates position of sRNA-*fliC* base pairing. (**B**) Immunoblot analysis showing UhpU and MotR overexpression leads to increased flagellin levels and FliX overexpression leads to reduced flagellin levels in the cytosol. Flagellin levels were determined by immunoblot analysis using α-FliC antibody. A sample from a Δ*fliC* strain was included as a control given the detection of a cross-reacting band slightly larger than flagellin. The Ponceau S-stained membrane serves as a loading control. Cells were grown with shaking at 180 rpm to OD_600_ ∼ 1.0, and cell fractions were separated by a series of centrifugation steps as detailed in Materials and Methods. (**C**) Northern blot analysis showing UhpU and MotR overexpression increases *fliC* mRNA levels and FliX overexpression reduces *fliC* levels across growth. The 5S RNA served as a loading control. The variation in *fliC* levels in the pBR* and pZE control samples is due to the different strain background (*crl^+^* versus *crl^-^*) and the length of membrane exposure. (**D**) Predicted base-pairing between *fliC* and UhpU, MotR or FliX. Seed sequences predicted by (Melamed *et al*., 2016) or by this study are underlined. Numbering is from AUG of *fliC* mRNA and +1 of indicated sRNAs. **Figure supplement 1.** Effects of UhpU, MotR* and FliX-S overexpression on flagellin and *fliC* mRNA levels. **Figure supplement 2.** *In vitro* structural probing of interaction between UhpU, MotR and FliX sRNAs with *fliC* mRNA.

We predicted base pairing between the three sRNAs and sequences overlapping the RIL-seq peaks internal to the *fliC* CDS (*Figure 4D*) and encompassing seed sequences suggested for the sRNAs (Melamed *et al*., 2016). To test for UhpU, MotR and FliX base pairing with these predicted sequences, we carried out *in vitro* footprinting with labeled fragments of the *fliC* mRNA (*Figure 4—figure supplement 2*). Upon cleavage with RNase III and lead, we observed changes in the regions predicted to be involved in base pairing (red brackets) that were dependent on the WT RNAs but not with derivatives carrying mutations in the regions predicted to be involved in base pairing. We also observed Hfq dependent changes (black bracket) in the region from ∼+40 to +66 from the *fliC* AUG, which is enriched for ARN motif sequences (AAA, AAT, AAC, AAG, AAC), known to be important for mRNA binding to the distal face of Hfq binding (reviewed in (Updegrove *et al*., 2016)). Additionally, we noted that both MotR and the MotR-M1 mutant RNAs led to additional protection at another region (thin red bracket) and increased cleavage (red asterisks) at other positions and suggesting a second region of MotR base pairing with *fliC* as well as MotR-induced structure changes. In general, the differences in cleavage by RNase III (preference for double-stranded RNA) and lead (preference for single-stranded RNA), indicate the *fliC* sequence from ∼+40 to ∼+170 is more structured than the surrounding regions. These differences in secondary structure could be the reasons for positive regulation by UhpU and MotR and negative regulation by FliX but also complicate analysis using standard reporter fusions with compensatory mutations.

### MotR and FliX modulate the S10 operon

Given that genes encoding ribosomal proteins were among the top MotR and FliX targets in the RIL-seq data sets and were not detected for many other sRNAs (*Supplementary file 1* and *Figure 3—figure supplement 1B*), we investigated MotR and FliX regulation of these genes. Several of the top interactions for MotR and FliX in the RIL-seq data mapped to the essential S10 operon, again within the CDSs (*Figure 5*, *Supplementary file 1*, and *Figure 3—figure supplement 1B*). The co-transcriptional regulation of the S10 operon has been studied extensively (Zengel & Lindahl, 1996, Zengel *et al*., 2002, Zengel & Lindahl, 1992). The leader sequence upstream of the first gene *rpsJ* encoding S10 is bound by the ribosomal protein L4, encoded by the third gene in the operon (*rplD*), causing transcription termination, thus modulating the levels of all the ribosomal proteins in the operon in response to the levels of unincorporated L4. L4 binding also has been shown to specifically inhibit translation of *rpsJ*, an effect that can be genetically distinguished from the L4 effect on transcription termination (Freedman *et al*., 1987).

**Figure 5.**
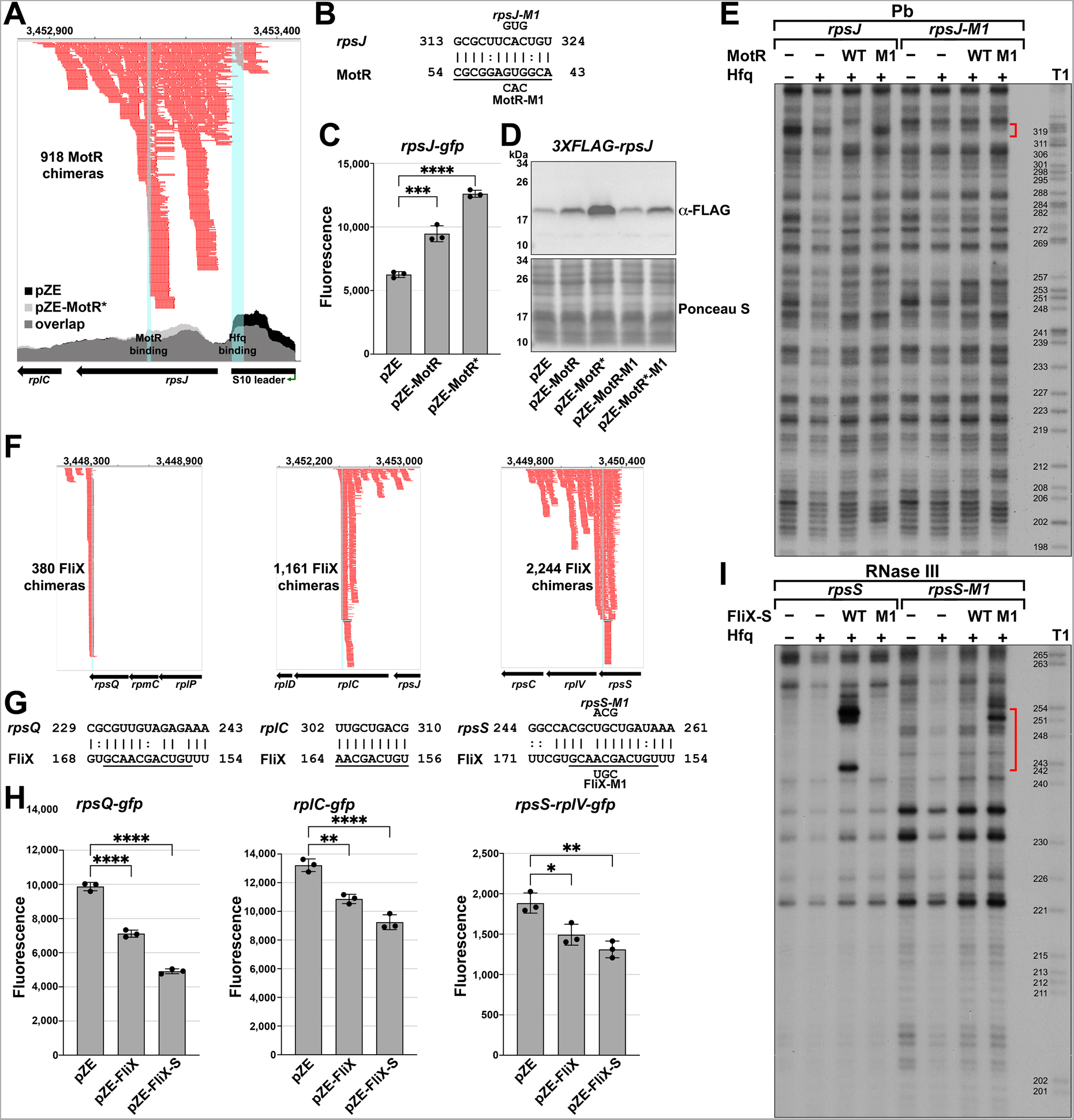
MotR and FliX base pair with the S10 mRNA leading to upregulation and downregulation, respectively. (A) Browser image showing MotR chimeras (in red) in S10 leader and *rpsJ* region. Data analyzed is from (RIL-seq experiment 1, (Melamed *et al*., 2020)). Coverage of the region in total RNA-seq libraries is shown for empty vector (pZE) and for pZE-MotR* overexpression (*Supplementary file 2*). The Hfq and MotR binding sites as detected in *Figure 5—figure supplement 2A* and *2B* are highlighted in light blue. (**B**) Base-pairing between *rpsJ* and MotR with sequences of mutants assayed. Predicted MotR seed sequence is underlined. Numbering is from +1 of *rpsJ* mRNA and MotR sRNA. (**C**) MotR induces *rpsJ-gfp* reporter fusion. Test of *rpsJ*-MotR interaction with reporter assays of *rpsJ-gfp* expressed from pXG10-SF with MotR or MotR* expressed from pZE. (**D**) MotR increases FLAG-tagged S10 levels. 3XFLAG-S10 was expressed from pBAD33 and MotR or MotR* was expressed from pZE. A mutation in MotR eliminates this regulation. 3XFLAG-S10 levels were determined by immunoblot analysis using α-FLAG antibody. The Ponceau S-stained membrane serves as a loading control. (**E**) RNase III-mediated cleavage of *rpsJ* directed by MotR in region of base pairing. ^32^P-labeled *rpsJ* and *rpsJ*-M1 were treated with lead for 10 min with or without MotR and MotR-M1 and separated on a sequencing gel. Region protected by MotR binding, which overlaps the predicted base pairing sequence, is indicated by the red bracket. Numbering is from +1 of *rpsJ* mRNA. (**F**) Browser image showing FliX chimeras (in red) in the S10 operon. Highlighted in light blue are the base pairing regions between FliX and the S10 operon mRNA. Data analyzed is from (RIL-seq experiment 1, (Melamed *et al*., 2020)). (**G**) Base pairing between *rplC, rpsS, rpsQ* and FliX with sequences of mutants assayed. FliX seed sequence predicted by (Melamed *et al*., 2016) is underlined. Numbering is from AUG of indicated CDS and +1 of FliX sRNA. (**H**) Test of FliX interactions with reporter assays of *rplC-gfp*, *rpsS-rplV-gfp* and *rpsQ-gfp* expressed from pXG10-SF or pXG30-SF with FliX or FliX-S expressed from pZE. (**I**) RNase III-mediated cleavage of *rpsS* directed by FliX-S in region of base pairing. ^32^P-labeled *rpsS* and *rpsS-M1* were treated with RNase III for 1.5 min with or without FliX-S and FliX-S -M1 and separated on a sequencing gel. Region protected by FliX binding, which overlaps the predicted base pairing sequence, is indicated by the red bracket. Numbering is from AUG of *rpsS* CDS. For (**C**) and (**H**), the average of three independent measurements is shown. Error bars represent one SD. One-way ANOVA comparison was performed to calculate the significance of the change in GFP signal (ns = not significant, * = P<0.05, ** = P<0.01, **** = P<0.0001). **Figure supplement 1.** Effects of MotR mutants on flagella number and *rpsJ* expression. **Figure supplement 2.** *In vitro* structural probing of interaction between MotR sRNA and *rpsJ* mRNA, and FliX sRNA and *rpsS* mRNA. **Figure supplement 3.** *In vivo* effects of FliX and FliX-S overproduction on *rpsS* mRNA.

To test for MotR regulation of *rpsJ* expression, we fused the S10 leader and part of the *rpsJ* CDS, including the position of the *rpsJ*-MotR chimeras (*Figure 5A*) and the region of predicted base-pairing (*Figure 5B*), to a GFP reporter (Corcoran *et al*., 2012, Urban & Vogel, 2009). MotR overexpression elevated the expression of the *rpsJ-gfp* fusion, and MotR* enhanced this effect (*Figure 5C*). Positive regulation of S10 expression by MotR and MotR* was similarly observed by immunoblot analysis of an N-terminal FLAG-tagged S10 protein encoded along with the S10 leader behind the heterologous promoter on a pBAD plasmid (*Figure 5D*). A mutation in the MotR seed sequence (MotR-M1 and MotR*-M1, *Figure 1—figure supplement 1A*) eliminated the up-regulation of the FLAG-tagged S10 (*Figure 5D*) and the MotR effect on flagella number (*Figure 5—figure supplement 1A*). To examine base pairing between MotR and the sequences internal to the *rpsJ* CDS, we carried out *in vitro* structure probing in the presence of Hfq (*Figure 5E* and *Figure 5—figure supplement 2A* and *2B*). The RNase T1, RNase III and lead cleavage assays supported the position of the predicted base-pairing between MotR and *rpsJ* mRNA, indicating MotR binds to *rpsJ* at ∼+150 nt in its CDS. Again, we detected Hfq binding (black bracket), here to the attenuator hairpin in the S10 leader sequence (*Figure 5—figure supplement 2B*), which has three ARN sequences (AGG, AGU and AAC). The M1 mutation eliminated binding in the predicted region of pairing but a complementary mutation in the corresponding region of *rpsJ* mRNA did not restore MotR binding (*Figure 5E*). We suggest that, as for the MotR target region of *fliC*, MotR binds to more than one site, the MotR target region of *rpsJ* is highly structured, and MotR and Hfq binding might all lead to conformational changes that compound the interpretation of the mutations.

Nevertheless, to further define the determinants needed for MotR-mediated up regulation, we generated a series of *rpsJ-gfp* fusions to include only the first seven amino acids of S10 removing the MotR base pairing site, to remove the S10 leader sequence, to remove stem D required for L4-mediated regulation, or to remove the attenuator hairpin stem E (*Figure 5— figure supplement 1B*). MotR-dependent regulation was eliminated for each of these constructs suggesting that S10 leader sequence is needed along with the MotR binding site internal to the *rpsJ* CDS for MotR-dependent regulation (*Figure 5—figure supplement 1B*). To test if Hfq binding to *rpsJ* is critical for the activation, we repeated the GFP reporter assay in an Hfq^Y25D^ mutant defective for binding ARN sequences on the distal face of the protein (Zhang *et al*., 2013a). Supporting a role for Hfq, MotR, which is present at the same levels in the Hfq WT and Hfq^Y25D^ mutant strains, no longer upregulates *rpsJ-gfp* in the distal face mutant background (*Figure 5—figure supplement 1C*). Collectively, our results are consistent with MotR base pairing internal to *rpsJ* affecting Hfq binding to the S10 leader sequence, which in turn results in increased *rpsJ* translation.

Based on the RIL-seq data, FliX interacts with multiple regions in the S10 operon mRNA, all internal to CDSs (*Figure 5F*). The predicted base-pairing regions (*Figure 5G*) align with the highest peaks of chimeras in the RIL-seq data and overlap with the seed sequence suggested for FliX (Melamed *et al*., 2016). We tested the effects of FliX on expression from this operon by constructing *gfp* fusions to regions of *rplC*, *rpsQ*, and *rpsS*-*rplV*. In all cases, overproduction of FliX or FliX-S led to a reduction in the expression of these fusions (*Figure 5H*). To test for a direct interaction between FliX-S and the *rpsS* mRNA, we again carried out structure probing (*Figure 5I* and *Figure 5—figure supplement 2C and 2D*). The regions that were changed in *rpsS* and FliX-S in the *in vitro* footprinting aligned with the predicted binding region between the two RNAs. Introduction of the M1 mutation (*Figure 1—figure supplement 1A*) eliminated FliX-S binding to the *rpsS* mRNA while introduction of a complementary mutation in the *rpsS* mRNA restored FliX-S-M1 binding (*Figure 5I*). We hypothesize that FliX downregulation of the *rplC*, *rpsQ*, and *rpsS*-*rplV* fusions as well as the *fliC* mRNA is due to sRNA-directed mRNA degradation. Further experiments are needed to test this model, but *in vivo* primer extension assays carried out for RNA isolated from in mid-log phase cells (OD_600_ ∼0.6) showed an increase in 5’ ends in proximity to the binding site on the *rpsS* mRNA in FliX or FliX-S overexpressing strains (*Figure 5—figure supplement 3*).

### Increased S10 levels correlate with increased readthrough of the flagellar operons

We wondered how the positive regulation of *rpsJ* by MotR might impact flagella synthesis. The S10 protein encoded by *rpsJ* has two roles in the cell. It is incorporated into the 30S ribosome subunit but also forms a transcription anti-termination complex with NusB (Lüttgen *et al*., 2002, Luo *et al*., 2008, Baniulyte *et al*., 2017). We evaluated the importance of each of the two S10 roles to flagella number by EM. First, we overexpressed a S10 mutant (S10Δloop) that is missing the ribosome binding loop but is still active in anti-termination (Luo *et al*., 2008) from an inducible plasmid and analyzed the number of flagella per cell. Cells carrying the S10Δloop plasmid had higher number of flagella like cells overexpressing MotR* (*Figure 6A*). We noted that overexpression of wild type S10 from the plasmid used for overexpression of S10Δloop did not lead to an increase in flagella number (*Figure 6A*), though presumably MotR is normally increasing flagella number by impacting the levels of the WT protein. Possibly, only a specific concentration of S10 relative to other ribosome proteins increases the S10 role as an anti-terminator. Since *rpsJ* is essential and cannot be deleted, we also examined the effect of MotR* overexpression in a Δ*nusB* strain that cannot form the S10-NusB anti-termination complex. In this background, the stimulatory effect of MotR* on flagella number was eliminated (*Figure 6B*) as is also observed for S10Δloop overexpression in the Δ*nusB* background (*Figure 6—figure supplement 1A*). Based on these observations, we hypothesize that increased S10 levels upon MotR overexpression leads to increased anti-termination of some of the long flagella operons.

**Figure 6.**
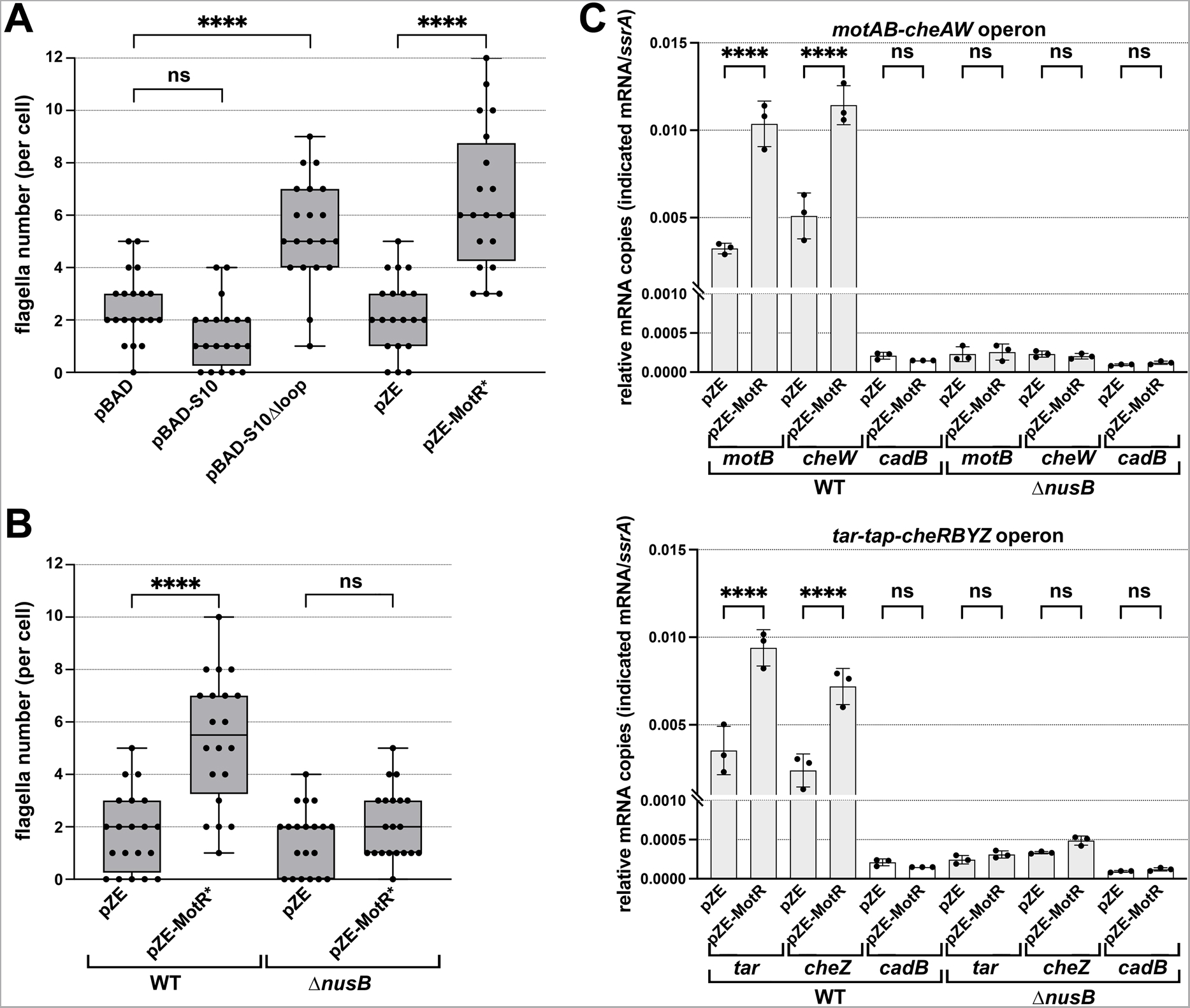
MotR overexpression leads to a *nusB*-dependent increase in expression from flagellar operons. (**A**) MotR* and S10Δloop overexpression increase the number of flagella. The number of flagella per cell detected by EM were counted for WT cells (GSO983) harboring the indicated plasmids. (**B**) MotR effect is eliminated in Δ*nusB* background. The number of flagella per cell detected by EM were counted for WT (GSO983) or Δ*nusB* cells (GSO1077) harboring the indicated plasmids. (**C**) MotR induces mRNA levels throughout the flagellar operons in WT background (GSO983) but not in Δ*nusB* background (GSO1077). MotR was expressed from pZE plasmid and the levels of *motB*, *cheW*, *tar*, *cheZ*, *ssrA* and *cadB* were monitored in comparison to their levels in the pZE control vector by RT-qPCR. *cadB* served as a non-flagellar gene control and *ssrA* served as a reference gene; the same *cadB* data is shown in both plots. Experiments were done in three biological replicates and one-way ANOVA comparison was performed to calculate the significance of the change in mRNA levels (ns = not significant, **** = P<0.0001). For (**A**) and (**B**), flagella were counted for 20 cells (black dots), and a one-way ANOVA comparison was performed to calculate the significance of the change in flagella number (ns = not significant, **** = P<0.0001). Box plot and error bars descriptions as in Figure 2. Each experiment was repeated three times and one representative experiment is shown. **Figure supplement 1.** Effects of MotR* and S10Δloop overexpression are lost in Δ*nusB* background.

To directly test this anti-termination hypothesis, we carried out RT-qPCR analysis in WT and Δ*nusB* backgrounds to examine the effects of MotR and MotR* overexpression on genes in the *motAB-cheAW* and *tar-tap-cheRBYZ* operons. For both operons, the mRNA levels of the tested genes were increased in WT upon MotR and MotR* overexpression (*Figure 6C* and *Figure 6—figure supplement 1B*). This increase was not observed for the non-flagellar control gene *cadB*. While the levels of the flagellar mRNAs in Δ*nusB* background were lower than in the WT, MotR and MotR* no longer induced these genes. Together these observations are consistent with the proposal that increased levels of non-ribosome associated S10 leads to increased levels of the S10-NusB anti-termination complex associated with RNA polymerase-σ^28^ and increased anti-termination of the long operons encoding flagellar proteins. It is also conceivable that even a slight upregulation of the S10 operon, as well as the S6 operon, given a significant number of MotR-*rpsF* chimeras (*Supplementary file 1*), along with anti-termination of *rrn* operons, could lead to more active ribosomes, which are needed for flagellar protein synthesis. On the other hand, a negative effect of FliX on ribosomal components, which could reduce the number of active ribosomes, would be consistent with the repressive role of this sRNA.

### MotR and FliX have opposing effects on the expression of middle and late flagella genes

In a parallel line of experimentation, we examined the impact of overexpressing MotR* and FliX on the transcriptome by RNA-seq analysis (*Supplementary file 2*). The transcripts whose levels increased most with MotR* overexpression compared to the vector control (*Figure 7A*) corresponded predominantly to late genes and, to a lesser extent, middle genes, of the flagellar regulon. Of the 332 genes whose expression increased significantly (FDR = 0.05) by MotR* overexpression, 40 are reduced significantly (FDR = 0.05) in a strain lacking σ^28^ (Δ*fliA*) (Fitzgerald *et al*., 2014) (*Figure 7—figure supplement 1A*). Additionally, the sequence motif found for the promoters of the transcription units for which expression increased the most (FDR = 0.05 and ≥2 fold) upon MotR* overproduction (*Figure 7—figure supplement 1B*) is nearly identical to a σ^28^ recognition motif (Fitzgerald *et al*., 2014, Shi *et al*., 2020). In contrast, transcripts for flagellar genes were reduced by FliX overexpression (*Figure 7B*). Specifically, 28 of 149 genes for which the expression is reduced significantly (FDR = 0.05) are middle or late genes of the flagellar regulon (Fitzgerald *et al*., 2014). We note that we did not observe differential levels of the S10 operon transcript in the RNA-seq analysis upon FliX overexpression but did detect decreased levels of some transcripts encoding ribosomal proteins upon MotR* overexpression (*Figure 7B* and *Supplementary file 2*). However, the total RNA for the RNA-seq experiments was isolated from cells early in growth (OD_600_ ∼0.2).

**Figure 7.**
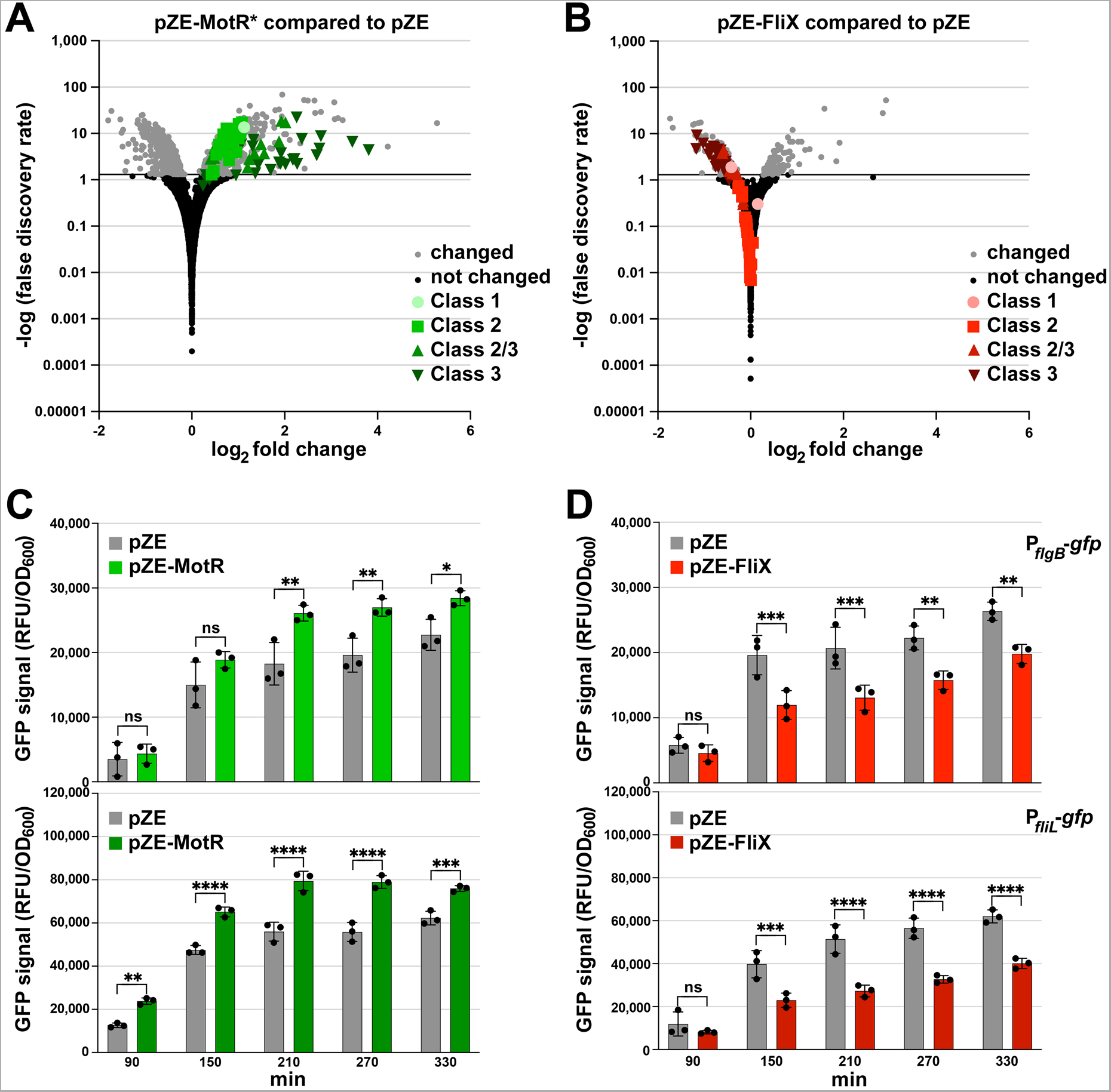
MotR and FliX overproduction leads to increased and decreased expression of flagellar genes, respectively. (**A**) MotR* induces flagellar genes. Green symbols represent flagellar regulon genes as indicated on the graph. (**B**) FliX reduces flagellar genes. Red symbols represent flagellar regulon genes as indicated on the graph. In (A) and (B) Differential expression analysis was conducted with DESeq2 and threshold for differentially expressed transcripts was set to adjusted value of p < 0.05. (**C**) MotR overexpression increases the activity of GFP fusions to P*_flgB_* and P*_fliL_*. The activity of the promoters was monitored for 330 min by measuring the GFP signal and dividing it with the culture OD_600nm_ in the presence of MotR expressed from pZE plasmid. (**D**) FliX overexpression decreases the activity of GFP fusions to P*_flgB_* and P*_fliL_*. The activity of the promoters was monitored for 330 min by measuring the GFP signal and dividing it with the culture OD_600nm_ in the presence of FliX expressed from pZE plasmid. For (**A**) and (**B**), WT (GSO983) harboring the control vector pZE or the MotR* or the FliX expression plasmid were grown to OD_600_ ∼ 0.2; total RNA was extracted and used for the construction of cDNA libraries, which were analyzed as described in Materials and Methods. For (**C**) and (**D**), three biological repeats are shown in the graph. One-way ANOVA comparison was performed to calculate the significance of the change in GFP signal (ns = not significant, * = P<0.05, ** = P<0.01, *** = P<0.001, **** = P<0.0001). The experiments presented in 7C and *Figure 7— figure supplement 2B*, and in 7D and *Figure 7—figure supplement 2A*, were carried out on same day, respectively, and the same pZE samples are shown. **Figure supplement 1.** Overlap in MotR* overexpression profile with σ^28^ regulon. **Figure supplement 2.** Effects of MotR* and FliX-S overexpression on P*_flgB_*-*gfp,* P*_fliL_*-*gfp* and FlgJ-SPA expression.

The effects of MotR, MotR*, FliX and FliX-S on flagella gene expression were further examined by monitoring fluorescence from *gfp* fused to the promoters of *flgB*, a representative Class 2 promoter, and *fliL*, a representative Class 2/3 promoter (Zaslaver *et al*., 2006). MotR and MotR* overexpression increased the activity of the two promoters, while FliX and FliX-S overexpression led to a reduction of their activity (*Figure 7C* and *7D*, *Figure 7—figure supplement 2A* and *2B*). The levels of C-terminally SPA-tagged FlgJ, also encoded by a Class 2 gene, similarly increased across growth upon MotR* overexpression, particularly early in growth, and decreased upon FliX-S overexpression (*Figure 7—figure supplement 2C* and *2D*). The data suggest that in addition to modulating anti-termination and/or ribosomal protein synthesis (*Figure 6*), MotR and FliX more broadly effect transcription initiation at flagellar genes though we do not know the mechanism. In general, these results are coherent with a positive effect of MotR on flagella synthesis and a negative effect of FliX.

### MotR increases and FliX decreases flagella synthesis

To examine the impact of chromosomally-encoded MotR and FliX on flagella synthesis and the flagellar regulon, we introduced the three-nucleotide M1 substitutions in the seed sequences of *motR* and *fliX* (MotR-M1 and FliX-M1, *Figure 1—figure supplement 1A*) at their endogenous chromosomal positions, avoiding the disruption of the nearby genes. MotR-M1 RNA levels were comparable to WT MotR levels (*Figure 8—figure supplement 1A*). The prominent ∼200 nt FliX band was reduced for FliX-M1, while other FliX processing products were affected less (*Figure 8—figure supplement 1B*).

We first examined the flagella number and motility for these strains. The *motR-M1* chromosomal mutation was associated with a moderate reduction in flagella number at two time points (OD_600_ ∼ 0.6 and 2.0) (*Figure 8A*), while slightly higher numbers of flagella were observed for the *fliX-M1* strain at the later time point (OD_600_ ∼ 2.0) (*Figure 8B*). In motility assays carried out as in *Figure 2*, we found reduced motility of the *motR-M1* strain compared to WT but no change was observed for the *fliX-M1* strain (*Figure 8—figure supplement 1C* and *1D*)). We also compared the motility of the *motR-M1* and *fliX-M1* strains to WT strains by mixing strains transformed with plasmids expressing either GFP or mCherry. WT strains expressing GFP were mixed with *motR-M1* or *fliX-M1* cells expressing mCherry or vice versa, and their motility was compared on 0.3% agar. For both combinations of WT and *motR-M1,* the fluorescent signal produced by the WT strain was more extensive than the fluorescent signal generated by *motR-M1* mutant outside of the site of inoculation (*Figure 8C*). Thus, in two independent assays, the *motR-M1* mutant exhibits reduced motility compared to the WT strain, while no significant difference was observed between WT and *fliX-M1* (*Figure 8D*).

**Figure 8.**
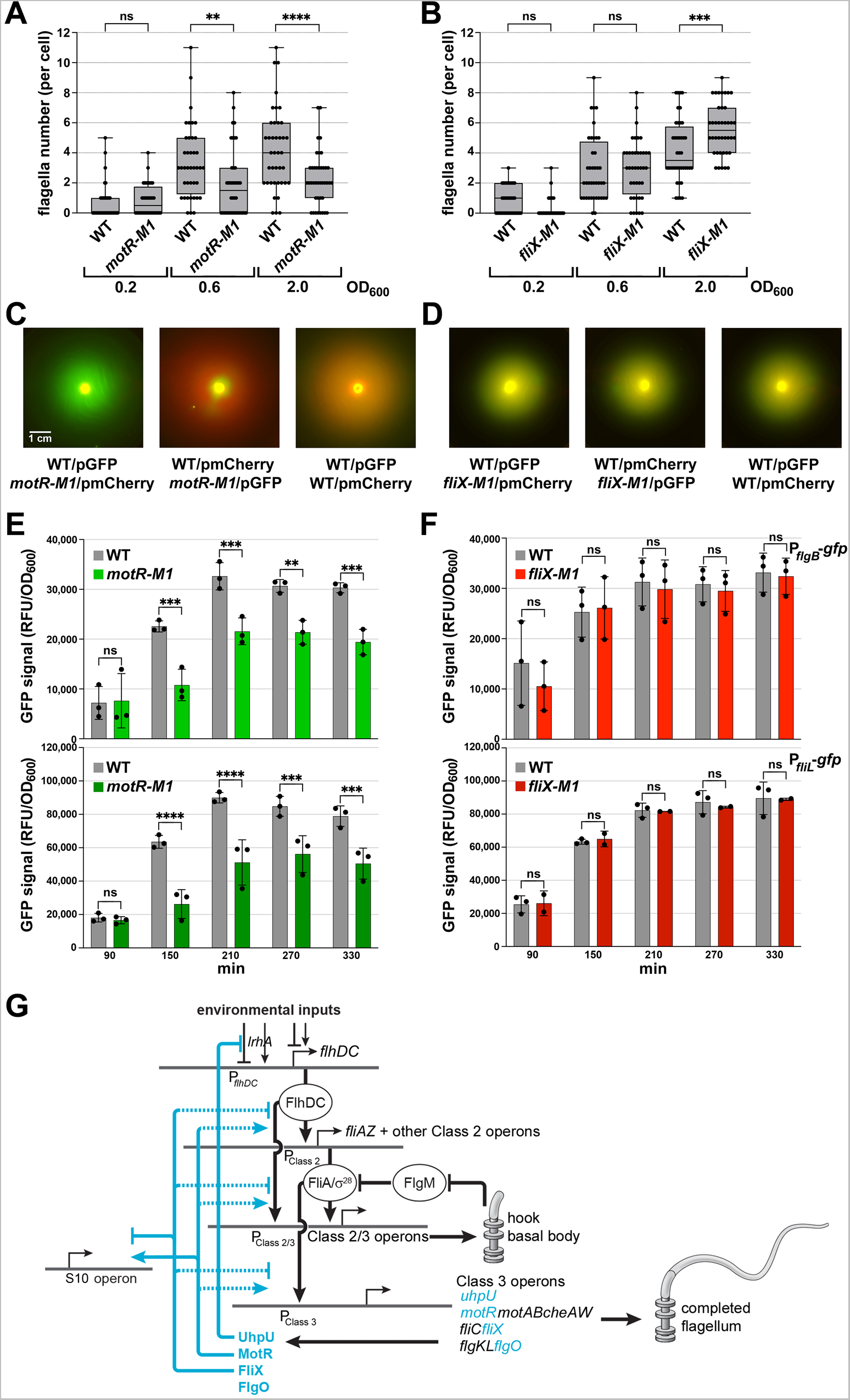
Complex regulatory network of sRNAs controlling flagella synthesis. (**A**) Reduction in flagella number in *motR-M1* mutant. (**B**) Increase in flagella number in *fliX-M1* mutant. (**C**) Reduced motility in *motR-M1* mutant (GSO1087) based on a competition assay with its corresponding WT (GSO1088). (**D**) No difference in motility in *fliX-M1* mutant (GSO1076) based on a competition assay with its corresponding WT (GSO983). (**E**) Reduction in P*_flgB_*-*gfp* and P*_fliL_*_-_*gfp* expression in *motR-M1* mutant (GSO1087) background compared to WT background (GSO1088). (**F**) No difference in P*_flgB_*-*gfp* and P*_fliL_*_-_*gfp* expression in *fliX-M1* mutant (GSO1076) background compared to WT background (GSO983). (**G**) σ^28^-dependent sRNAs control flagella synthesis at different levels. UhpU activates the flagellar regulon by repressing a regulator of *flhDC*. MotR and FliX, respectively, activate and repress middle and the late gene expression (dotted line indicates exact mechanism is not known, though we document base pairing with the *fliC* mRNA). MotR and FliX also connect ribosome and flagella synthesis by regulating genes in the S10 operon (solid line indicates documented base pairing with this mRNA). In (**A**) and (**B**), the number of flagella per cell detected by EM were counted for 40 cells (black dots) for the *motR-M1* (GSO1087) and its corresponding WT (GSO1088), and for *fliX-M1* (GSO1076) and its corresponding WT (GSO983), strains at three points in growth (OD_600_ ∼ 0.2, OD_600_ ∼ 0.6, and OD_600_ ∼ 2.0). A one-way ANOVA comparison was performed to calculate the significance of the change in flagella number (ns = not significant, ** = P<0.01, *** = P<0.001, **** = P<0.0001). Each experiment was repeated three times and one representative experiment is shown. Box plot and error bars descriptions as in Figure 2. For (**C**) and (**D**), WT or the corresponding mutant, expressed either green fluorescent signal or red fluorescent signal by carrying pCON1.proC-GFP or pCON1.proC-mCherry plasmid, respectively. In the left images, WT cells expressing GFP were mixed with mutant cells expressing mCherry; in the middle images, WT cells expressing mCherry were mixed with mutant cells expressing GFP; in the right images, WT cells expressing GFP were mixed with WT cells expressing mCherry. The indicated mixed cultures were spotted on a soft agar (0.3%) plate, incubated at 30°C, and imaged after 18 h. For (**E**) and (**F**), three biological repeats are shown in the graph (except for P*_fliL_*_-_*gfp* in *fliX-M1*, for which two repeats are shown). One-way ANOVA comparison was performed to calculate the significance of the change in GFP signal (ns = not significant, ** = P<0.01, *** = P<0.001, **** = P<0.0001). The scale given in (**C**) is the same for all motility plates, respectively. **Figure supplement 1.** Effects of chromosomal *motR-M1* and *fliX-M1* mutations. **Figure supplement 2.** Conservation of σ^28^-dependent sRNAs.

We also assessed the effects of the chromosomal mutations on the *flgB-gfp* and *fliL-gfp* fusions (Figure 8) as well as on FlgJ-SPA and *fliC* mRNA levels (*Figure 8—figure supplement 1*). The *motR-M1* mutant showed reduced activity of the two promoters (*Figure 8E*), in line with the increased activity of the promoters that was observed upon MotR overexpression (*Figure 7C*). The *fliX-M1* mutant showed similar activity of the two promoters in comparison to WT (*Figure 8F*). In western and northern analyses of the *motR-M1* strain compared to its parental WT, a delayed initiation of FlgJ-SPA and *fliC* mRNA synthesis in the mutant was observed, respectively (*Figure 8—figure supplement 1E* and *1G*). In contrast, FlgJ-SPA and *fliC* mRNA levels in the *fliX-M1* strain increased compared to the parental WT strain (*Figure 8—figure supplement 1F* and *1H*).

While negative effects of the *motR-M1* mutation on flagella number, motility, and flagellar gene expression were observed in all assays, positive effects of the *fliX-M1* mutation were only detected for flagella number, FlgJ-SPA protein, and *fliC* mRNA levels. However, for both sRNAs the mutation phenotype is opposite that of the overexpression phenotype. Collectively these observations indicate that MotR, expressed earlier in growth, increases flagella synthesis by positively regulating the middle and the late genes, while FliX, whose levels peak later, decreases flagella synthesis by downregulating the flagellar regulon. Thus, MotR and FliX, along with UhpU, add another layer of regulation to the flagellar regulon (*Figure 8G*).

## Discussion

In this study, we describe four *E. coli* sRNAs whose expression is dependent on σ^28^. We found three of these sRNAs affect flagella number and bacterial motility. Although previous studies showed that base pairing sRNAs act on the *flhDC* mRNA (Thomason *et al*., 2012, De Lay & Gottesman, 2012), our results revealed that the effect of sRNAs on flagellar synthesis is far more pervasive. Intriguingly, two of the σ^28^-dependent sRNAs show opposite effects. MotR, expressed earlier in growth, increases expression of flagellar and ribosomal proteins along with flagella number, while FliX, expressed later in growth, decreases expression of the proteins and flagella number. Thus, the two sRNAs, respectively, might be considered an accelerator and a decelerator for flagellar synthesis.

### Non-canonical mechanisms of sRNA action

Most commonly, sRNAs base pair with the 5’ UTRs of mRNA targets or at the very beginning of the CDS, primarily affecting ribosome binding or mRNA stability. However, MotR and FliX bind in the middle or even close to the ends of their target CDSs in the *fliC* gene and S10 operon. For both *fliC* and the S10 operon, the consequences of MotR and FliX overexpression are different. MotR leads to higher levels of *fliC* and the S10 protein, whereas FliX leads to lower levels of *fliC* and three genes in the S10 operon. We suggest that the positive and negative regulatory effects of MotR and FliX, respectively, occur by the same mechanisms on the *fliC* and S10 transcripts, with MotR changing the conformation of the RNAs and FliX leading to increased cleavage. However, these suggested mechanisms needed to be investigated further in future experiments. It is also noteworthy that, based on RIL-seq data, more examples of this type of interaction remain to be characterized.

Given that our study made extensive use of RIL-seq data, it provides an opportunity to evaluate these data. While RIL-seq provides a comprehensive map of RNA-RNA interactions that take place on Hfq under a specific condition, some caution about the interpretation is warranted as the interactions represent multiple types of relationships between two RNAs. As was found by a recent study (Faigenbaum-Romm *et al*., 2020), we suggest that if an interaction is highly abundant and discovered under multiple conditions, the sRNA is more likely to have a regulatory impact on the target mRNA though the mechanisms may be unknown. We noticed that the spread of the RIL-seq signal varies significantly between targets. One possible explanation for multiple peaks and a broad distribution is more than one base pairing site for the sRNA on the mRNA, but this hypothesis requires further testing. We predict additional studies of sRNA-target pairs with different types of RIL-seq signals will give further insights into the mechanisms and outcomes of base pairing.

The most studied and conserved sRNA-binding protein in gram-negative bacteria is Hfq. However, there are other sRNA-binding proteins (reviewed in (Melamed, 2020)). Among these is ProQ, which was shown to have overlapping, complementary, and competing roles with Hfq in *E. coli* (Melamed *et al*., 2020). Interestingly, ProQ was found to affect motility and chemotaxis in *S. enterica* (Westermann *et al*., 2019). In the absence of ProQ, the target sets for the σ^28^-dependent sRNAs on Hfq were changed significantly in *E. coli* (Table S5 in (Melamed *et al*., 2020)) suggesting that competition between Hfq and ProQ for binding RNAs likely also influences this regulatory circuit. In this context, it is worth noting that FlgO, the fourth σ^28^-dependent sRNA, which originates from the 3’UTR of the *flgL* and strongly binds Hfq (Melamed *et al*., 2020), does not have many targets. Possibly, FlgO has a role in titrating Hfq from other sRNAs or proteins, or in the recruitment of other proteins to a complex with Hfq. Interestingly, while the overall sequence of *flgO* is conserved in other bacterial species (*Figure 8—figure supplement 2*), the nucleotides in one of the single stranded loops (*Figure 1—figure supplement 1*) differ in *S. typhimurium*, possibly suggesting distinct regulatory mechanisms in different bacteria.

### Conservation of σ^28^-dependent sRNAs

We were surprised to find so many σ^28^-dependent Hfq-binding sRNAs and wondered about their phylogenic distribution. A previous study assessing the conservation of the *motR* and *uhpU* promoters showed that, while the *motR* promoter is well conserved across proteobacteria species, the *uhpU* promoter was not, implying different evolutionary pressures (Fitzgerald *et al*., 2018). Interestingly, however, a sRNA named RsaG, which originates from the 3’ UTR of *uhpT* and also is induced by glucose-6-phosphate, was found in the Gram-positive bacterium, *Staphylococcus aureus* (Bronesky *et al*., 2019). Although there is no sequence similarity between UhpU and RsaG, and RsaG has not been reported to regulate flagella synthesis, the independent evolution of regulatory sRNAs at the 3’ UTRs of *uhpT* in two disparate bacterial species is intriguing. RsaG was found to regulate redox homeostasis and to adjust metabolism to changing environmental conditions (Desgranges *et al*., 2022). While we focused on the UhpU role in the flagellar regulon and in controlling motility, the sRNA has many targets that are part of different metabolic pathways and redox homeostasis (*Supplementary file 1*), hinting at parallels between the two sRNAs.

The σ^28^-dependent sRNAs themselves are conserved among some of the Enterobacteriaceae (*Figure 8—figure supplement 2*) and thus may play a role in pathogenicity. During the preparation of this manuscript, two studies describing the application of RIL-seq to *S. enterica* and Enteropathogenic *E. coli* Hfq were published (Mizrahi *et al*., 2021, Matera *et al*., 2022). The RIL-seq analyses were carried out for cells grown under conditions that do not favor flagellar gene expression, but UhpU, MotR, FliX and FlgO were detected, confirming their synthesis and association with Hfq in pathogenic bacteria. It is also likely that still other sRNA regulators of the flagellar regulon remain to be characterized. In *S. enterica*, a leader RNA originating from the *mgtCBR* virulence operon was shown to affect the synthesis of one of the two flagellin genes that exist in this bacterium, impacting virulence and motility (Choi *et al*., 2017). In neonatal meningitis-causing *E. coli*, a sRNA missing from the *E. coli* MG1655 strain used in our study, was shown to reduce *fliC* mRNA levels (Sun *et al*., 2022).

### Roles of σ^28^-dependent sRNAs

The UhpU RIL-seq target set includes many flagellar regulon genes and some transcription regulators of the flagellar regulon, such as LrhA (Lehnen *et al*., 2002), hinting at a mechanism by which UhpU can affect flagella number and bacterial motility. However, since UhpU can also be derived from the *uhpT* mRNA (*Figure 1—figure supplement 1B*) and is predicted to have many targets that participate in carbon and nutrient metabolism (*Supplementary file 1*), we suggest this sRNA may play a broader role in linking carbon metabolism with flagella synthesis and motility.

MotR and FliX each have more limited target sets in the RIL-seq data but may comprise a unique regulatory toggle. While the transcription of the two sRNAs is dependent on the same sigma factor and they base pair in the CDS of targets in the same operons, base pairing results in opposing regulation. MotR, which is transcribed from within *flhC* at the top of the flagellar regulon, reaches its highest levels earlier than FliX and increases the flagella synthesis. In contrast, FliX, which is cleaved from the mRNA required to make the last protein needed to complete the flagellum, reaches its highest levels later in growth and appears to decrease flagella synthesis.

It is not yet clear how MotR and FliX base pairing with only a few targets can have such pervasive effects on flagellar gene expression and flagella number, but we suggest multiple mechanisms may be involved. One possibility is that the levels of flagellin encoded by *fliC*, up and down regulated by MotR and FliX, respectively, could be part of an autoregulatory loop that impacts the transcription of *flhDC* or other middle or late flagellar gene promoters. The increased and decreased levels of ribosomal proteins brought about by MotR and FliX regulation of the S10 operon also could impact the levels of available ribosomes, where even slight changes could have consequences given the high ribosome cost of flagella synthesis. Finally, we hypothesize that elevated levels of the S10 protein, due to the regulation by MotR, could, in conjunction with NusB, lead to increased anti-termination of long flagellar operons.

Based on our hypothesis that the MotR-mediated increase in S10 levels leads to increased anti-termination, we speculate that MotR activation of S10 expression could serve an autoregulatory role. Early in growth, transcription initiating from the σ^28^-dependent promoter in *flhC* terminates at the 5’ of *motA* generating MotR. As MotR levels increase, there is a concomitant increase in S10 levels, which could promote readthrough of the *motR* terminator leading to decreased MotR levels and increased full-length *motRAB-cheAW* mRNA.

In general, the σ^28^-dependent sRNAs add a new layer of regulation to the flagellar regulon and reinforce the conclusion that flagella synthesis is exquisitely regulated. The regulon will continue to serve as a model of a temporal and environmentally-controlled regulatory network with contributions from both transcription factors and regulatory RNAs.

## Materials and methods

### Key resources table

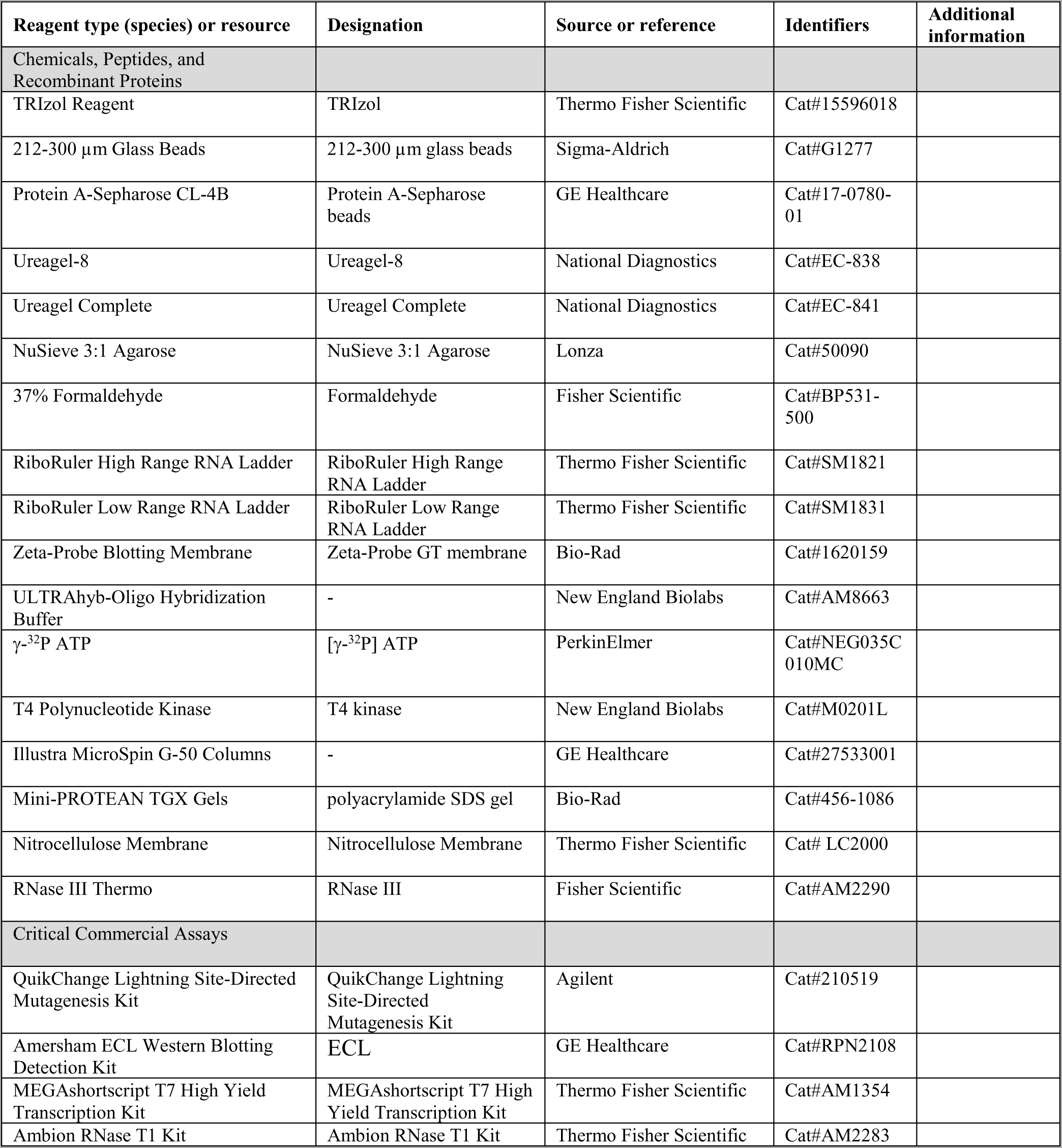
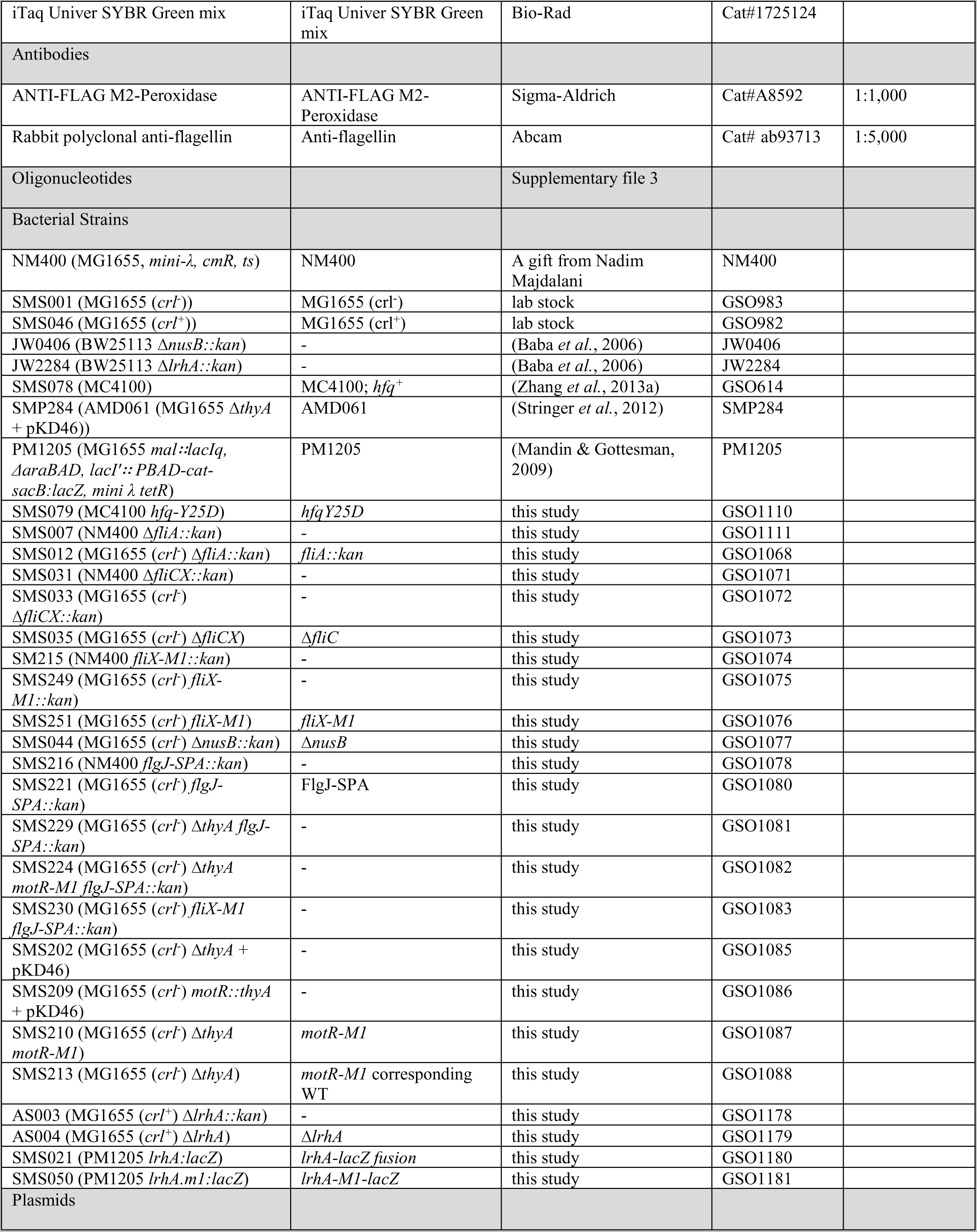
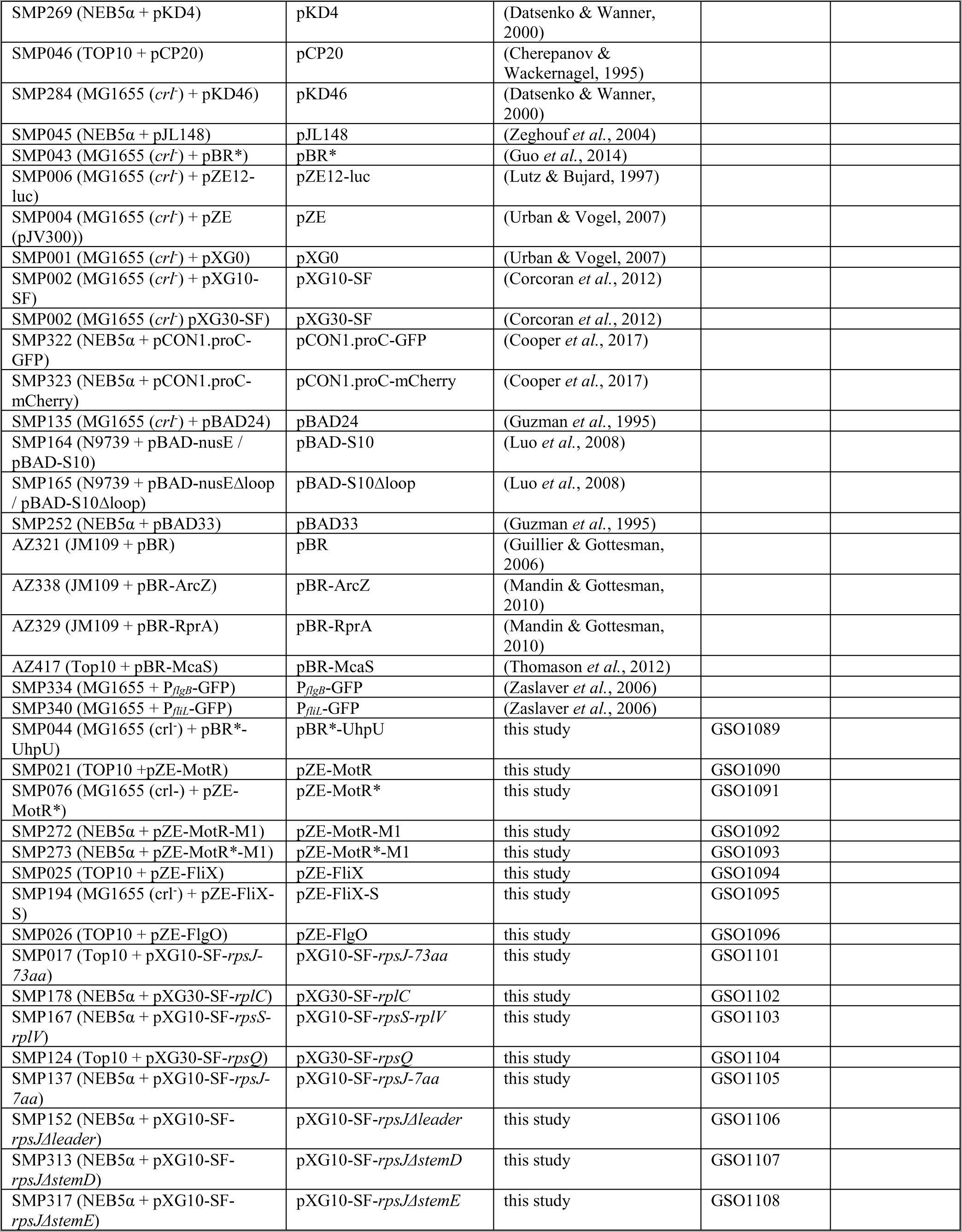
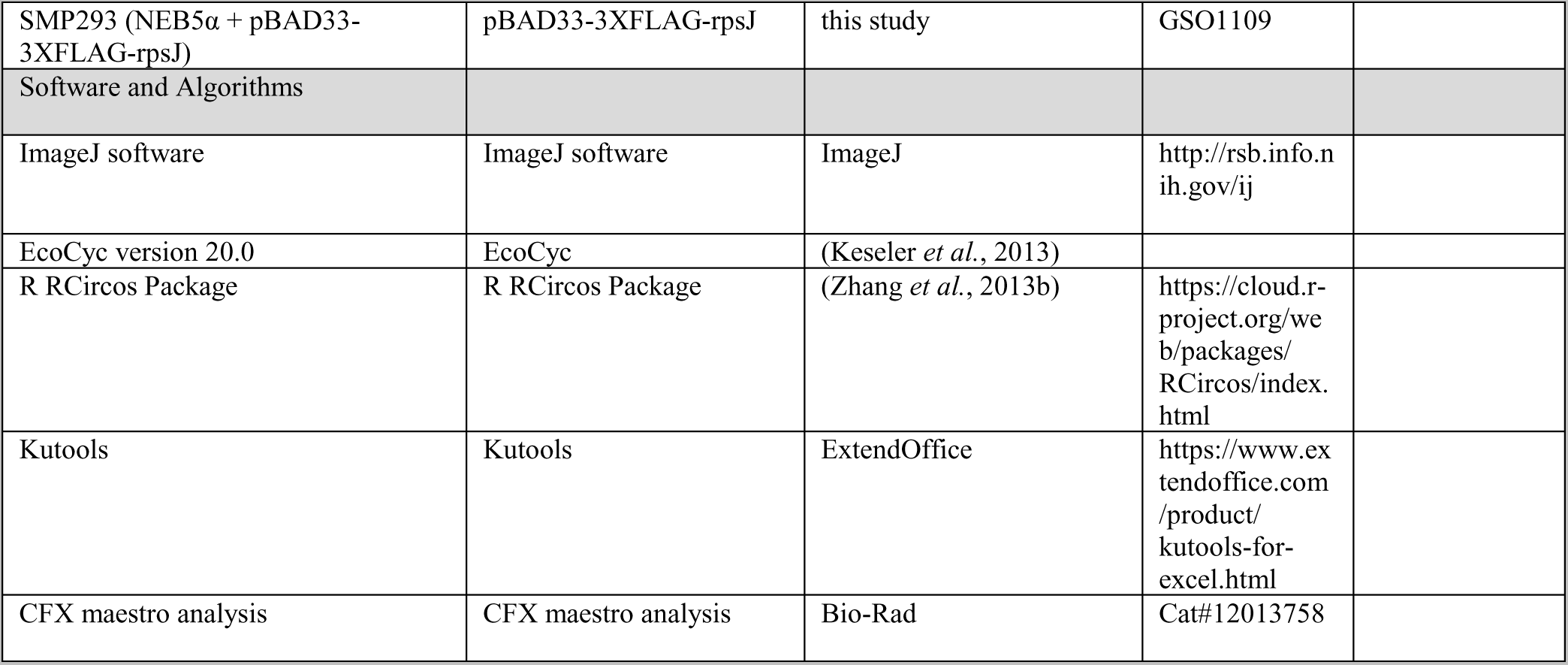

### Bacterial strains and growth conditions

*E. coli* MG1655 (GSO982 or GSO983) or MC4100 (GSO614) strains served as the WT strains in this study. All other bacterial strains studied here are listed in the *Key Resources Table* along with plasmids and oligonucleotides used. *E. coli* K-12 MG1655 genome was used as template to amplify mRNAs and sRNAs to be cloned into their respective sites. Unless indicated otherwise, all strains were grown with shaking at 250 rpm at 37°C in LB rich medium. Ampicillin (100 µg/ml), chloramphenicol (25 µg/ml), kanamycin (30 µg/ml), arabinose (0.2%) and IPTG (1 mM) were added where appropriate. Unless indicated otherwise, overnight cultures were diluted to an OD_600_ = 0.05 and grown for the indicated times or to the desired optical densities.

### Strain construction

*fliA::kan*, *fliCX::kan*, and *fliX-M1:kan* strains were constructed by amplifying the *kan^R^* sequence from pKD4 (Datsenko & Wanner, 2000) using oligonucleotides listed in *Supplementary file 3* and recombining (Datsenko & Wanner, 2000) the product into the chromosome of strain NM400 (kind gift of Nadim Majdalani). *flgJ* was SPA-tagged by amplifying the SPA sequence adjacent to *kan^R^*sequence from pJL148 (Zeghouf *et al*., 2004) using oligonucleotides listed in *Supplementary file 3* and recombining (Datsenko & Wanner, 2000) the product into the chromosome of strain NM400. *motR-M1* strain was constructed using the scar-free system, FRUIT (Stringer *et al*., 2012) as previously described. Briefly, *thyA* was deleted from MG1655 (*crl^-^*) (GSO983) strain by PCR amplification of Δ*thyA* from AMD061 (Stringer *et al*., 2012) followed by recombination using pKD46 (Datsenko & Wanner, 2000). Next, *thyA* was inserted back to the genome next to the site of mutation and selection was made by growth on minimal media lacking thymine. The *motR-M1* mutation was introduced while simultaneously removing *thyA*. The selection for colonies missing *thyA* was carried out using minimal medium M9 plates supplied with 0.4% glucose, 0.2% casamino acids, 20 µg/ml trimethoprim, and 100 µg/ml thymine. *lrhA::kan*, and *nusB::kan* deletion strains were obtained from other groups (Baba *et al*., 2006) as referenced in *Key Resources Table*. All deletions and mutations were confirmed by sequencing and then transferred to new backgrounds by P1 transduction. Where indicted, *kan^R^*was removed from the chromosome using plasmid pCP20 (Cherepanov & Wackernagel, 1995).

Construction of strains carrying chromosomal *lacZ* fusions was carried out using PM1205 as previously described (Mandin & Gottesman, 2009). In brief, the *lrhA* fragment was amplified using KAPA Hifi (Fisher Scientific) using oligonucleotides SM079 and SM080 (*Supplementary file 3*) and transformed into PM1205 with a series of selective screens on minimal media plates supplemented with sucrose, LB, LB supplemented with chloramphenicol, and LB supplemented with tetracycline. Mutagenesis of *lrhA-lacZ* fusion was achieved by recombineering an *lrhA-M1* sequence instead of the WT *lrhA* sequence, using gBlock listed in *Supplementary file 3*.

### Plasmid construction

Descriptions of plasmids used in this study are in *Supplementary file 3*. Construction of the constitutive overexpression plasmids was done according to (Urban and Vogel, 2009) using pZE12-luc. The IPTG-inducible UhpU overexpression plasmid was constructed using a pBRplac derivative harboring *kan^R^*, pMSG14 (Guo *et al*., 2014). The *uhpU* sequence, starting from its second nt, was amplified by PCR using oligonucleotides TU558 and TU561 (*Supplementary file 3*), digested with *Aat*II and *Hin*dIII and cloned into pMSG14 digested with the same restriction enzymes. 3XFLAG-*rpsJ* was expressed from pBAD33 (Guzman *et al*., 1995). The S10 leader and *rpsJ* sequence along with the 3XFLAG sequence was PCR amplified using oligonucleotides SM533 and SM435, digested with *Kpn*I and *Hin*dIII and cloned into pBAD33 digested with the same restriction enzymes. Construction of GFP-fusion plasmids was carried out principally as described in (Urban and Vogel, 2009), using the pXG10-SF or pXG30-SF (Corcoran et al., 2012). Briefly, regions of target genes, mainly regions captured in the chimeric fragments, were PCR amplified, digested with *Mph*1103I and *Nhe*I and cloned into pXG10-SF or pXG30-SF digested with the same restriction enzymes. The full list of oligonucleotides used in this study can be found in *Supplementary file 3*. Mutagenesis of the different plasmids was achieved using the QuikChange Lightning Site-Directed Mutagenesis Kit (Agilent). All plasmids were freshly transformed into the appropriate strains before each of the experiments.

### RNA isolation

Cells corresponding to the equivalent of 10-20 OD_600_ were collected, washed once with 1X PBS, and frozen in liquid nitrogen. RNA was extracted according to the standard TRIzol protocol (Thermo Fisher Scientific) as described previously (Melamed *et al*., 2020). At the last step, RNA was resuspended in 20-50 µl of DEPC water and quantified using a NanoDrop (Thermo Fisher Scientific).

### RNA coimmunoprecipitation (Co-IP) assay

RNAs that co-IP using polyclonal antibodies to Hfq were isolated as described (Zhang *et al*., 2002) with the following modifications. MG1655 (GSO983) was grown to OD_600_ ∼0.6 and ∼1.0 in LB medium. Cells corresponding to the equivalent of 20 OD_600_ were collected, and cell lysates were prepared by vortexing with 212-300 µm glass beads (Sigma-Aldrich) in a final volume of 1 ml of lysis buffer (20 mM Tris-HCl/pH 8.0, 150 mM KCl, 1 mM MgCl_2_, 1 mM DTT). Co-IPs were carried out using 100 ml of α-Hfq, 120 mg of protein A-Sepharose beads (GE Healthcare), and 950 µl of cell lysate. Co-IP RNA was isolated from protein A-Sepharose beads by extraction with phenol: chloroform:isoamyl alcohol (25:24:1), followed by ethanol precipitation. Total RNA was isolated from 50 ml of cell lysate by TRIzol (Thermo Fisher Scientific) extraction followed by chloroform extraction and isopropanol precipitation. Total and co-IP RNA samples were resuspended in 15 µl of DEPC H_2_O, and 5 µg total RNA and 0.5 µg co-IP RNA were subjected to northern analysis as described below.

### Northern blot analysis

For smaller RNAs, total RNA (5 μg) was separated on a denaturing 8% polyacrylamide urea gel containing 6 M urea (1:4 mix of Ureagel Complete to Ureagel-8 (National Diagnostics) with 0.08% ammonium persulfate) in 1X TBE buffer at 300V for 90 min. The RNA was transferred to a Zeta-Probe GT membrane (Bio-Rad) at 20 V for 16 h in 0.5X TBE. For longer RNAs, total RNA (10 μg) were fractionated on formaldehyde-MOPS agarose gels as previously described (Adams *et al*., 2017). Briefly, RNA was denatured in 3.7% formaldehyde (Fisher), 1X MOPS (20 mM MOPS, 5 mM NaOAc, 1 mM EDTA, pH 7.0) and 1X RNA loading dye (Thermo Fisher Scientific) for 10 min at 70°C and incubated on ice. The RNA was loaded onto a 2% NuSieve 3:1 agarose (Lonza), 1X MOPS, 2% formaldehyde gel and separated at 125-150V at 4°C for 1-2 h and then transferred to a Zeta-Probe GT membrane (Bio-Rad) via capillary action overnight (Streit *et al*., 2009). For both types of blots, the RNA was crosslinked to the membranes by UV irradiation. RiboRuler High Range and Low Range RNA ladders (Thermo Fisher Scientific) were marked by UV-shadowing. Oligonucleotide probes (listed in *Supplementary file 3*) for the different RNAs were labelled with 0.3 mCi of [γ-^32^P] ATP (Perkin Elmer) by incubating with 10 U of T4 polynucleotide kinase (New England Biolabs) at 37°C for 1 h.

### Primer extension assay

Primer extension analysis was performed using an oligonucleotide (listed in *Supplementary file 3*) specific to the *rpsS* as described (Zhang *et al*., 1998). RNA samples (5 µg of total RNA) were incubated with 2 pmol of 5-^32^P-end-labeled oligonucleotide primer at 80°C and then slow-cooled to 42°C. After the addition of dNTPs (1 mM each) and AMV reverse transcriptase (10 U, Life Sciences Advanced Technologies Inc.), the reactions were incubated in a 10-μl reaction volume at 42°C for 1 h. The reactions were terminated by adding 10 μl of Stop Loading Buffer. The cDNA products then were fractionated on 8% polyacrylamide urea gels containing 8 M urea in 1X TBE buffer at 70 W for 70 min.

### RT-qPCR

Total RNA was isolated from cultures grown to OD_600_∼0.2 and RNA concentrations were determined using a NanoDrop. Samples were treated with DNase using TURBO DNA-free™ Kit (Thermo Fisher Scientific). DNA-free RNA was used for cDNA synthesis using iScript cDNA Synthesis Kit (Bio-Rad) and cDNA concentrations were measured by Qubit fluorimeter (Invitrogen). Equal amounts of cDNA were loaded into 96-well plate and cDNA was quantified by CFX Connect Real-Time system (Bio-Rad) using iTaq Univer SYBR Green mix (Bio-Rad) according to manufacturer instructions. Specific oligonucleotide primers were designed for each gene and the expression was normalized using *ssrA* levels. Serial dilutions of *E. coli* genomic DNA in known concentrations were used to generate a standard curve. Starting quantity of cDNA samples were determined based on the standard curve and normalization was done using the starting quantities of *ssrA*. Cq was measured in duplicate or triplicate for each biological sample. CFX maestro analysis software was also used for conducting the analysis.

### RNA structure probing

GeneBlock fragments carrying the *motR*, *fliX*, *rpsJ* or *rpsS* CDS (IDT) were used as DNA templates for *in vitro* transcription with MEGAshortscript T7 High Yield Transcription Kit (Invitrogen). The transcripts were dephosphorylated with calf intestinal alkaline phosphatase (CIP, New England Biolabs) and then radioactively labeled at 5’ end with [γ-^32^P] ATP (Perkin Elmer) and T4 kinase (Invitrogen), and purified on an 8% polyacrylamide/7M urea gel and eluted in buffer containing 20 mM Tris-HCl/pH 7.5, 0.5 M NaOAc, 10 mM EDTA and 1% SDS at 4°C for overnight, followed by ethanol precipitation. The RNA concentration was determined by measuring the OD_260_ on Nanodrop.

For all the structural probing assays, 0.2 pmole of the labeled transcript, 2 pmole of unlabeled transcript and 1 µg of yeast RNA with or without 2 pmole (hexameric concentration) of purified Hfq were mixed in 10 µl of 1x Structural Buffer in Ambion RNase T1 Kit (Invitrogen). The reactions were incubated at 37°C for 10 min, followed by treatment at 37°C with 0.02 U RNase T1 for 10 min, 1.3 U RNase III for 1.5 min, or 50 µmole lead acetate for 10 min, whereupon 20 µl Inactivation Buffer and 1 µl Glycoblue were added. The RNAs were precipitated and resuspended in 10 µl Loading Buffer, and analyzed on a 8% polyacrylamide/7 M urea gel run in 1x TBE. RNase T1 and alkali digestion ladders of the end-labeled transcripts were used as molecular size markers.

### Translational reporter assay

The GFP reporter assays were carried out essentially as described (Melamed et al., 2016). Overnight cultures were grown in 2 ml of LB media supplemented with the appropriate antibiotics at 37°C with constant shaking at 250 rpm. Cells were then diluted to OD_600_∼0.05 in 1 ml of fresh LB medium supplemented with the appropriate antibiotics in 96-well plate and grown at 37°C with constant shaking at 250 rpm for 3 h. Cells were pelleted and resuspended in filtered 1 X PBS. Fluorescence was measured using the BD LSRFortessa or Beckman Coulter Cytoflex flow cytometer. The level of regulation was calculated by subtracting the auto-fluorescence and then calculating the ratio between the fluorescence signal of a strain carrying the sRNA over-expressing plasmid and the signal of a strain carrying the control plasmid. Three biological repeats were prepared for every sample.

The β-galactosidase assays were carried out as described (Miller, 1992). Overnight cultures grown as for the GFP reporter assays were diluted 1:100 into 5 ml of fresh LB with antibiotic and 0.2% arabinose and grown at 37°C with constant shaking at 250 rpm until OD_600_ ∼ 0.7. IPTG (1 mM) was added to cells harboring inducible sRNAs plasmids. After β-galactosidase activity was measured, the Miller units were calculated from the following formula:

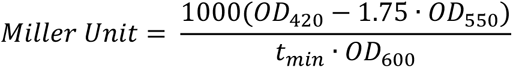

### Transcriptional reporter assays

Overnight cultures harboring *flgB-gfp* and *fliL-gfp* fusions (Zaslaver *et al*., 2006) were grown as described for the translation reporter assays and then diluted to OD_600_∼0.05 in 150 µl of fresh LB medium supplemented with the appropriate antibiotics in a transparent bottom 96-well plate. Bacterial growth and promoter activity were monitored for 330 min at 37°C using OD_600_ and GFP fluorescent measurements, respectively, using a Synergy H1 plate reader (Agilent).

### Immunoblot analysis

Bacteria were grown to the desired OD_600_, and the cells in 0.5 ml – 4 ml of culture were collected. Cell lysates were prepared by resuspending cell pellets with SDS-loading buffer normalized to the cell density, and samples were then heated for 10 min at 95°C. Protein samples were subjected to a 4%-15% polyacrylamide SDS gel electrophoresis followed by electrotransfer to a nitrocellulose membrane (Fisher Scientific). The membrane was blocked with 3% milk, probed with anti-flagellin antibodies (1/10,000) (Abcam) and then with anti-rabbit secondary antibody (1/10,000) or with ANTI-FLAG M2-Peroxidase (HRP) (1/1,000), (Sigma-Aldrich). Signals were visualized by the ECL system (Biorad).

### Flagellin measurements

WT (GSO983) cells harboring pBR*, pBR*-UhpU, pZE, pZE-MotR, pZE-MotR*, pZE-FliX or pZE-FliX-S were grown with shaking at 180 rpm in 5 ml of LB at 37°C to OD_600_ ∼ 1.0. Cell pellets collected by centrifugation were suspended in 5 ml of PBS and then heated at 65°C for 5 min, followed by centrifugation to obtain the cell pellets and supernatants, which contained the cytoplasmic flagellin molecules and depolymerized flagellin monomers, respectively. The cell pellets were resuspended in the SDS-loading buffer, normalized to the cell density. Proteins in the supernatants were precipitated by 10% trichloroacetic acid, resuspended in Tris/SDS loading buffer and heated at 95°C for 10 min.

### Electron microscopy

Overnight cultures were diluted in fresh medium and grown with shaking at 180 rpm, at 37°C to mid-log phase (OD_600_∼0.6-0.8) unless indicated otherwise. Cells were collected by centrifugation at 1,000 rpm for 20 min, and pellet was resuspended in 300 µl of saline. Next, 3 µl of bacterial suspension were placed on a freshly glow-discharged carbon covered electron microscopic support grid (EMS, Hatfield, PA) for 5 min. The grid was washed twice with distilled water and stained for 1 min with 0.75% aqueous solution of uranyl formate, pH 4.5. The grids were imaged in Thermo Fisher Scientific (Hillsboro, OR) FEI Tecnai 20 electron microscope operated at 120 kV. The images were recorded using AMT (Woburn, MA) XR81 CCD camera. Flagella were counted for 20-40 cells in each sample as indicated in the Figure legends. Each analysis was repeated a minimum of three times.

### Motility assays

Overnight cultures (∼1 µl) were spotted onto 0.3% soft agar plates by touching the agar softly with the tip and ejecting the culture. Plates were incubated right-side up at 30°C above a beaker filled with water for 9-24 h. Plates were made with the appropriate antibiotics and with 1 mM IPTG. The plates were imaged using Bio-Rad imager (using Colorimetric settings) and the diameter of the bacterial culture was calculated using the ImageJ software. Two technical repeats and three biological repeats were carried out for each strain. For motility competition assays cells were first transformed with pCON1.proC-GFP or pCON1.proC-mCherry plasmids (Cooper *et al*., 2017), resulting in a GFP or an mCherry signal, respectively. In each case, equal numbers of bacterial cells based on OD_600_ of each overnight culture for one strain expressing a green fluorescence signal and a second strain expressing a red fluorescent signal were mixed before spotting them onto 0.3% soft agar plate and the plates were incubated as described above. Images were taken using Bio-Rad imager with the following settings: Colorimetric (1-2 sec) for bright field, Cy2 for GFP (auto optimal exposure), Cy3 for mCherry (auto optimal exposure). Images were merged using Image Lab (Bio-Rad).

### RNA-seq

Overnight cultures were diluted in fresh LB medium and grown to early-log phase (OD_600_∼0.2). RNA was extracted using the standard TRIzol protocol (Thermo Fisher Scientific) as described above. Total RNA libraries were constructed using the RNAtag-Seq protocol with a few modifications to allow capture of short RNA fragments as previously described (Melamed *et al*., 2018). The libraries were sequenced by paired-end sequencing using the HiSeq 2500 system (Illumina) at Molecular Genomics Core, *Eunice Kennedy Shriver* National Institute of Child Health and Human Development. RNA-seq data processing follows the same procedures as RIL-seq data analysis for QC analysis, adaptor removal, and alignment with the Python RILSeq package (Melamed *et al*., 2018). The raw fastq records were demultiplexed with python script index_splitter.py (https://github.com/asafpr/RNAseq_scripts/blob/master/index_splitter.py) followed by adapter removal with cutadpt software (version 3.4). The trimmed fastq reads were mapped to the *E. coli* genome (ecoli-k12-MG1655-NC_000913-3) with Python RILSeq package (version 0.74, https://github.com/asafpr/RILseq). Deeptools software (version 3.5.1) was used to generate bigwig file for coverage visualization. Read counts were obtained with featureCounts tool of Subread software (version 2.0.3) and a customized annotation file based on EcoCyc version 20.0 (Keseler *et al*., 2013) with manual addition of sRNAs and small proteins from (Hör *et al*., 2020). Differential expression analyses were conducted with R DESeq2 package (Love *et al*., 2014) and default normalization. Deferentially expressed genes were extracted with the parameter of ‘independentFiltering=FALSE’. The sequencing data reported in this paper have been deposited in GEO under accession number GSE1774487.

### Determination of sequence motifs and base-pairing predictions

Common binding motifs were searched with MEME software (Bailey *et al*., 2009). Genes that were induced the most by MotR* overexpression in RNA-seq (*Supplementary file 2*) (FDR = 0.05 and ≥2 fold) were extracted from the data and divided to transcription units based on EcoCyc version 20.0 (Keseler et al., 2013). For each transcription unit, genomic sequence was extracted using coordinates for the start codon of the first gene in the transcription unit and 250 nt upstream to the gene. For sRNAs, genomic sequence was extracted using coordinates for the transcription start site and 250 nt upstream to the gene. For outputs, motif length was restricted to 28.

Base-pairing regions between two RNAs were predicted using IntaRNA (Mann *et al*., 2017) or TargetRNA2 (Kery *et al*., 2014).

### Functional annotation analysis

Functional annotation analysis of sRNAs targets was carried out using the Database for Annotation, Visualization and Integrated Discovery (DAVID) (Huang da *et al*., 2009). Gene names served as the input list in each case. Targets that were present in at least three RIL-seq conditions in Table S1 were included in the analysis.

### Circos plots

Circos plots follow the procedures of R RCircos Package (Zhang *et al*., 2013b). Link lines are used to label the statistically significant chimeric fragments (S-chimeras as defined in (Melamed *et al*., 2016)). RIL-seq from six different growth conditions was analyzed and S-chimeras present in at least four of the six conditions are included in the plots.

### Browser images

Data from RIL-seq experiment 1 from Melamed at al., 2020 extracted for unified S-chimeras files for the different sRNAs were mapped based on the first nt of each read in the chimera. BED files are generated with Python RILSeq package (Melamed *et al*., 2018) and viewed using the UCSC genome browser (Kent *et al*., 2002). For previously annotated RNA in GTF file, BED files are directly generated with command of generate_BED_file_of_endpoints.py and EcoCyc ID. For genes annotated in the current study, significant chimeras which involve the relevant gene are first extracted from significant interaction file, then chimeric reads involving the S-chimeras are extracted from chimeric read file. To be a qualified chimeric read, RNA1 start position of the read must overlap with the genomic range of RNA1 in S-chimera and RNA2 start position of the read must overlap with the genomic range of RNA2 in S-chimera. Finally, the read list for genes annotated in the current study is supplied to generate_BED_file_of_endpoints.py command to generate BED file.

## Supporting information

Melamed-Supplementary file 1

Melamed-Supplementary file 2

Melamed-Supplementary file 3

## Acknowledgements

We thank M. Gottesman for plasmids expressing wild type and *rpsJ* mutants, O. Steele-Mortimer for plasmids constitutively expressing GFP or mCherry, and D. Court for the S10 antibody. We thank J. Wade for sharing the sequences used to generate the σ^28^ binding motif and J. Wade and G. Baniulyte for advice on the FRUIT method. We thank the NICHD Molecular Genomics Core, particularly Tianwei Li, for all the library sequencing. We also appreciate the help of A. Peer with the sRNA conservation analysis. We are grateful to the Storz and S. Gottesman labs for all the helpful discussions and thank the Storz lab, S. Gottesman, and J. Wade for their comments on the manuscript. This work utilized the computational resources of the NIH HPC Beowulf cluster (http://hpc.nih.gov).

## Funding

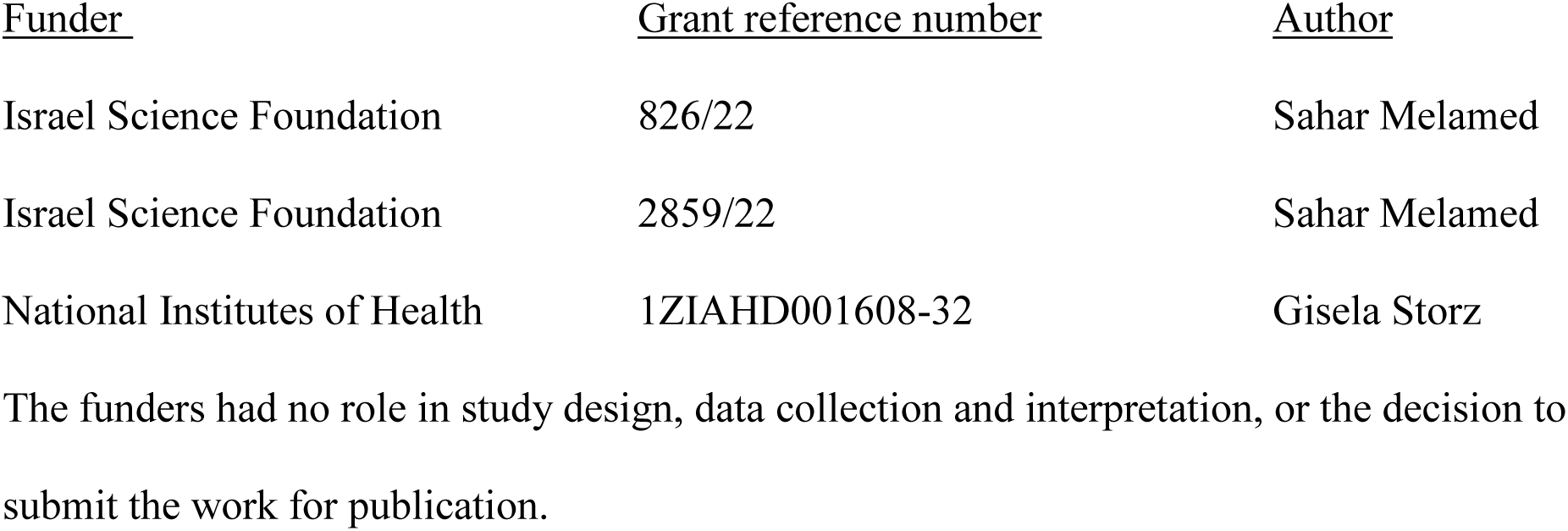

## Author contributions

S.M. and G.S. conceived of the project. S.M., A.Z., J.M., and A.S. designed, performed, and analyzed the experiments. M.J. performed all EM. H.Z. performed all computational analyses under the supervision of S.M.. S.M., A.Z. and G.S. prepared the figures and wrote the manuscript. G.S. supervised the project.

## Figure Supplement Legends

**Figure 1—figure supplement 1.**
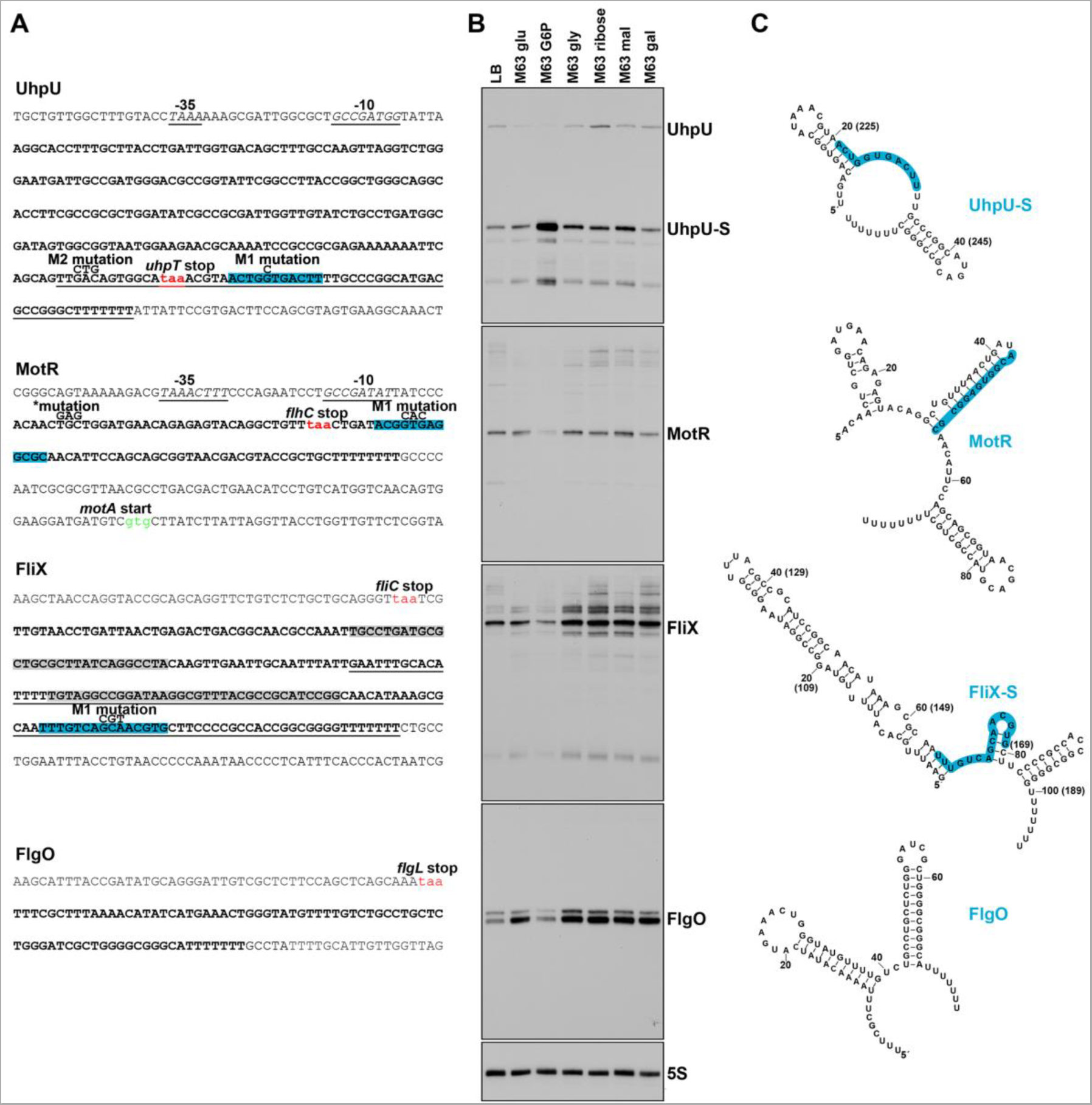
Sequences and predicted structures of UhpU, MotR, FliX and FlgO sRNAs and effect of carbon source on sRNA levels. (**A**) Genomic regions of σ^28^-dependent sRNAs. sRNAs sequences are in bold. The two FlgO bands (*Figure 1C and 1D*) likely result from differential processing and the bold sequence corresponds to the higher molecular weight band. Stop codons are indicated in red whereas start codons are indicated in green. The -10 and - 35 regions of the *uhpU* and *motR* promoters are in italics and underlined. Regions highlighted in blue reflect the suggested seed sequence of *uhpU* (Melamed *et al*., 2016) and the regions that base pair with the targets tested in this study for *motR* and *fliX*. The *uhpU-S* and *fliX-S* sequences are underlined. The *uhpU-M1*, *uhpU-M2*, *motR\**, *motR-M1* and *fliX-M1* mutations are labeled in the *uhpU*, *motR* and *fliX* sequences, respectively. The regions highlighted in gray denote the REP sequences in *fliX*. (**B**) Expression of σ^28^-dependent sRNAs in cells grown in different carbon sources. Total RNA was extracted from WT (GSO983) grown to exponential phase (OD_600_∼0.6) in LB medium or M63 minimal medium supplemented with 0.2% of glucose, glucose-6-phosphate (G6P), ribose, maltose, or galactose or 0.4% glycerol, separated on an acrylamide gel and sequentially probed for UhpU, MotR, FliX, FlgO and 5S RNAs. This is the same membrane probed for the RbsZ sRNA and 5S RNA (same panel) in Supp. Figure S8A of (Melamed *et al*., 2020). (**C**) Structures of UhpU-S, MotR, FliX-S and FlgO predicted by mfold (Zuker, 2003). Numbering for full-length UhpU and FliX is indicated in parentheses. Base pairing regions are highlight in blue as in (**A**).

**Figure 1—figure supplement 2.**
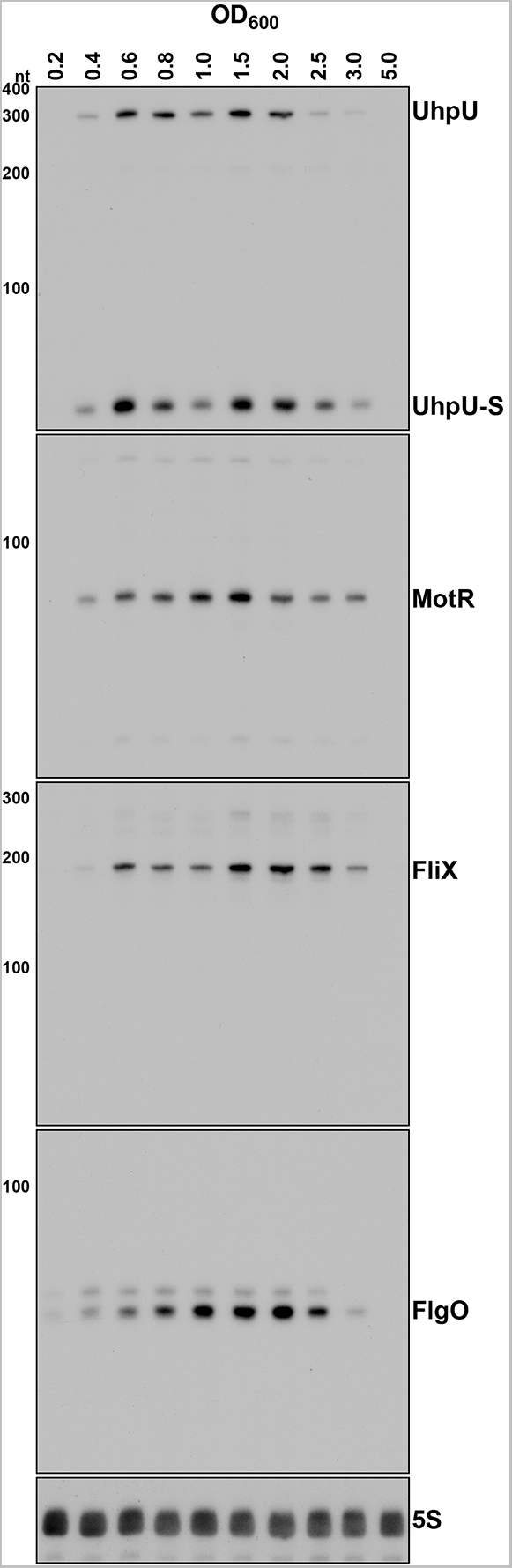
UhpU, MotR, FliX and FlgO levels across growth. Expression of σ^28^-dependent sRNAs in cells across growth. Total RNA was extracted from WT (GSO983) cells grown to the indicated time points. RNA was separated on an acrylamide gel and sequentially probed for all four σ^28^-dependent sRNAs and the 5S RNA. A full-length transcript (∼260 nt) and several processed transcripts, of which one is predominant (UhpU-S, ∼60 nt), are detected for UhpU, one prominent band (∼95 nt) is detected for MotR, one prominent band (∼200 nt) is detected for FliX, and two close bands close in size (∼75 nt) are detected for FlgO.

**Figure 2—figure supplement 1.**
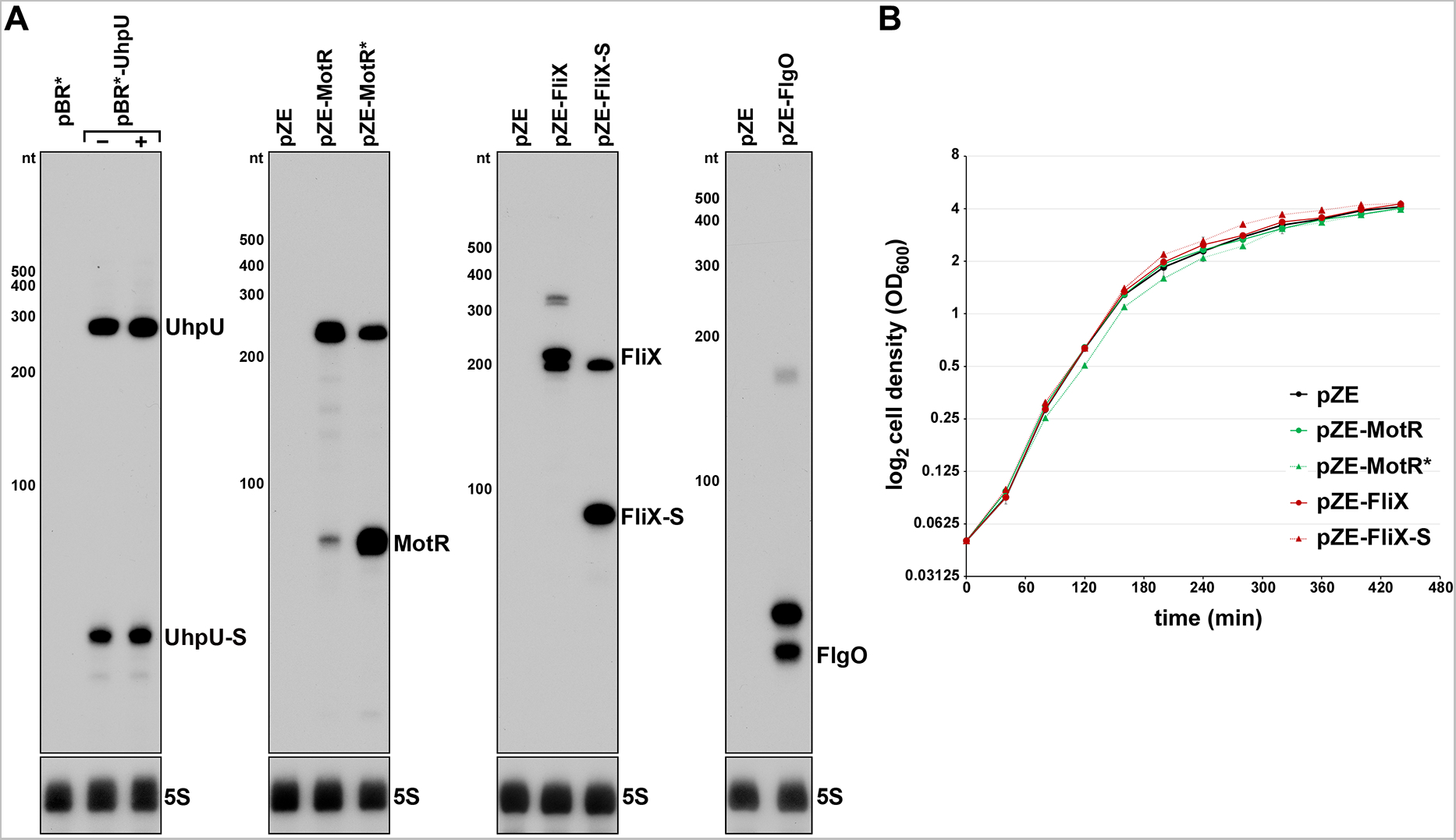
Expression of σ^28^-dependent sRNAs from plasmids and the effect of MotR and FliX overexpression on growth. (**A**) Expression of σ^28^-dependent sRNAs from plasmids. Total RNA was extracted from WT (GSO983), harboring a plasmid as indicated in the figure, grown to exponential phase (OD_600_∼0.2) with or without 1 mM IPTG. RNA was separated on an acrylamide gel and sequentially probed for an sRNA (UhpU, MotR, FliX or FlgO) and 5S RNAs. Higher bands were detected for MotR and FliX expressed from the pZE plasmid, likely due to some terminator readthrough. (**B**) Growth curves of WT (GSO983), harboring a plasmid as indicated in the figure, in LB medium. Overnight cultures were diluted to OD_600_ = 0.05 and cell growth was monitored for 440 min by OD_600_ measurements.

**Figure 2—figure supplement 2.**
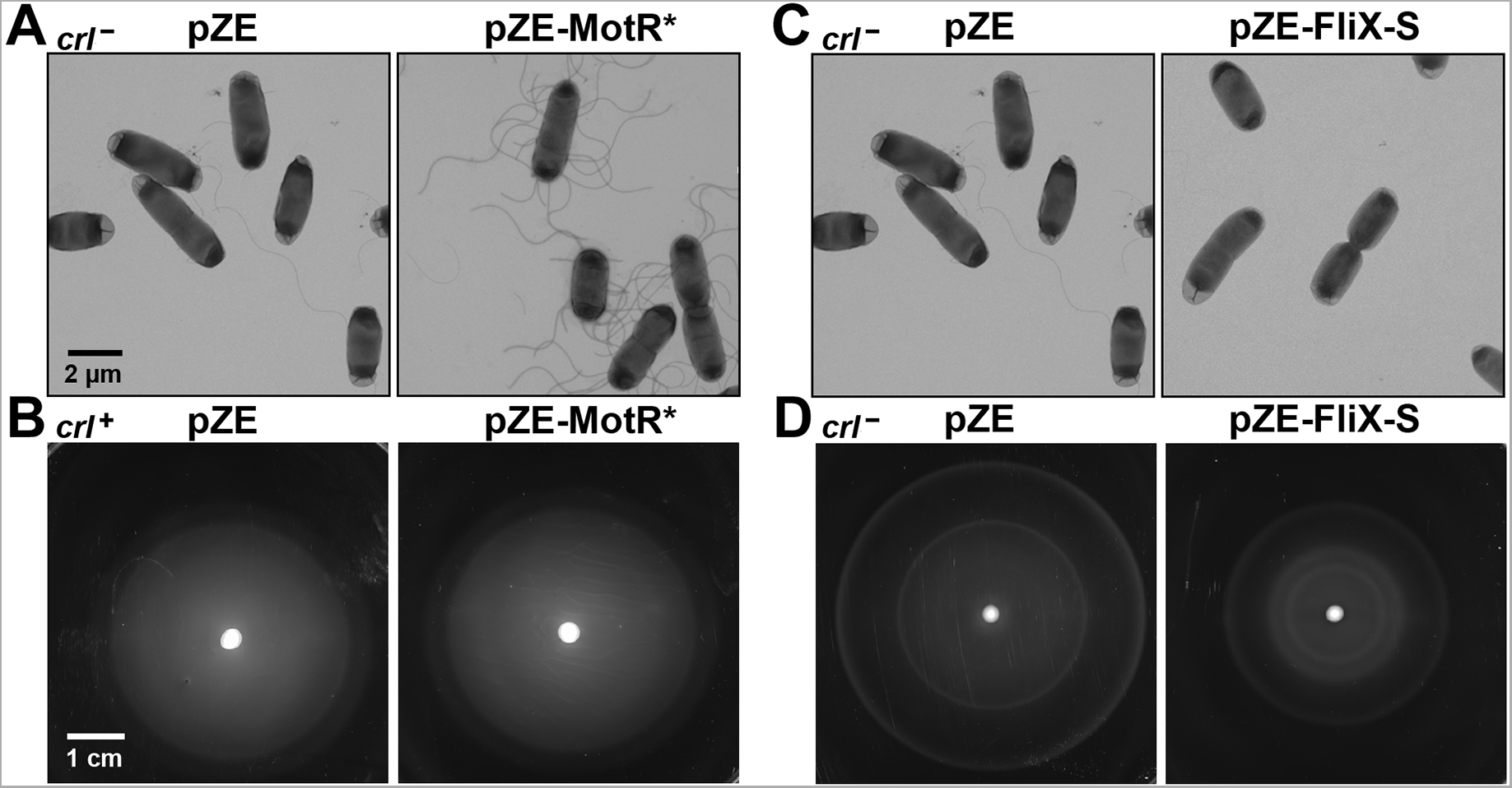
Effects of MotR* and FliX-S overexpression on flagella number and motility. (**A**) Increase in flagella number with MotR* overexpression based on EM analysis for WT (*crl*^-^) cells carrying an empty vector or overexpressing MotR*. (**B**) No significant change in motility with MotR* overexpression based on motility in 0.3% agar for WT (*crl*^+^) cells carrying an empty vector or overexpressing MotR*. (**C**) Reduction in flagella number with FliX-S overexpression based on EM analysis for WT (*crl*^-^) cells carrying an empty vector or overexpressing FliX-S. (**D**) Reduced motility with FliX-S overexpression based on motility in 0.3% agar for WT (*crl*^-^) cells carrying an empty vector or overexpressing FliX-S. The scales given in (**A**) and (**B**) are the same for all EM images and all motility plates, respectively.

**Figure 3—figure supplement 1.**
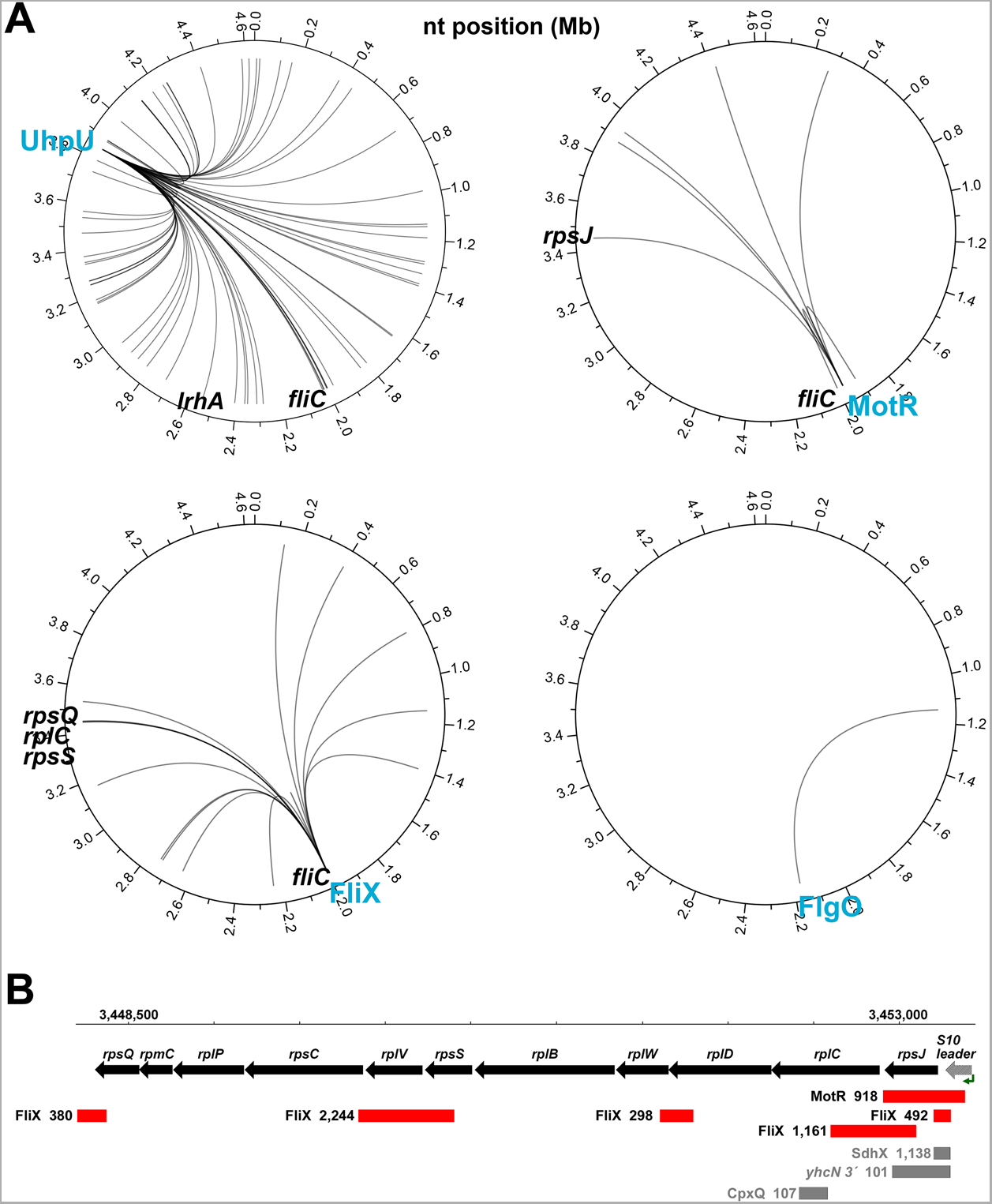
Interactomes for σ^28^-dependent sRNAs. (**A**) Circos plots showing σ^28^-dependent sRNAs targets that were detected in at least four of the six RIL-seq conditions. Each sRNA has a unique pattern of partners. UhpU is hub that binds hundreds of targets, MotR and FliX have smaller datasets and FlgO only has one target even though it is one of the most abundant sRNAs on Hfq. Targets that were addressed in the paper are labeled on the plots. Circos plots were drawn using R RCircos Package (Zhang *et al*., 2013b). (**B**) Schematic representation of the S10 operon with positions of sRNA binding. sRNAs that have more than 100 chimeras with genes from the S10 operon are shown. Numbers in the brackets represent the number of chimeras. Data analyzed is from (RIL-seq experiment 1, (Melamed *et al*., 2020)).

**Figure 4—figure supplement 1.**
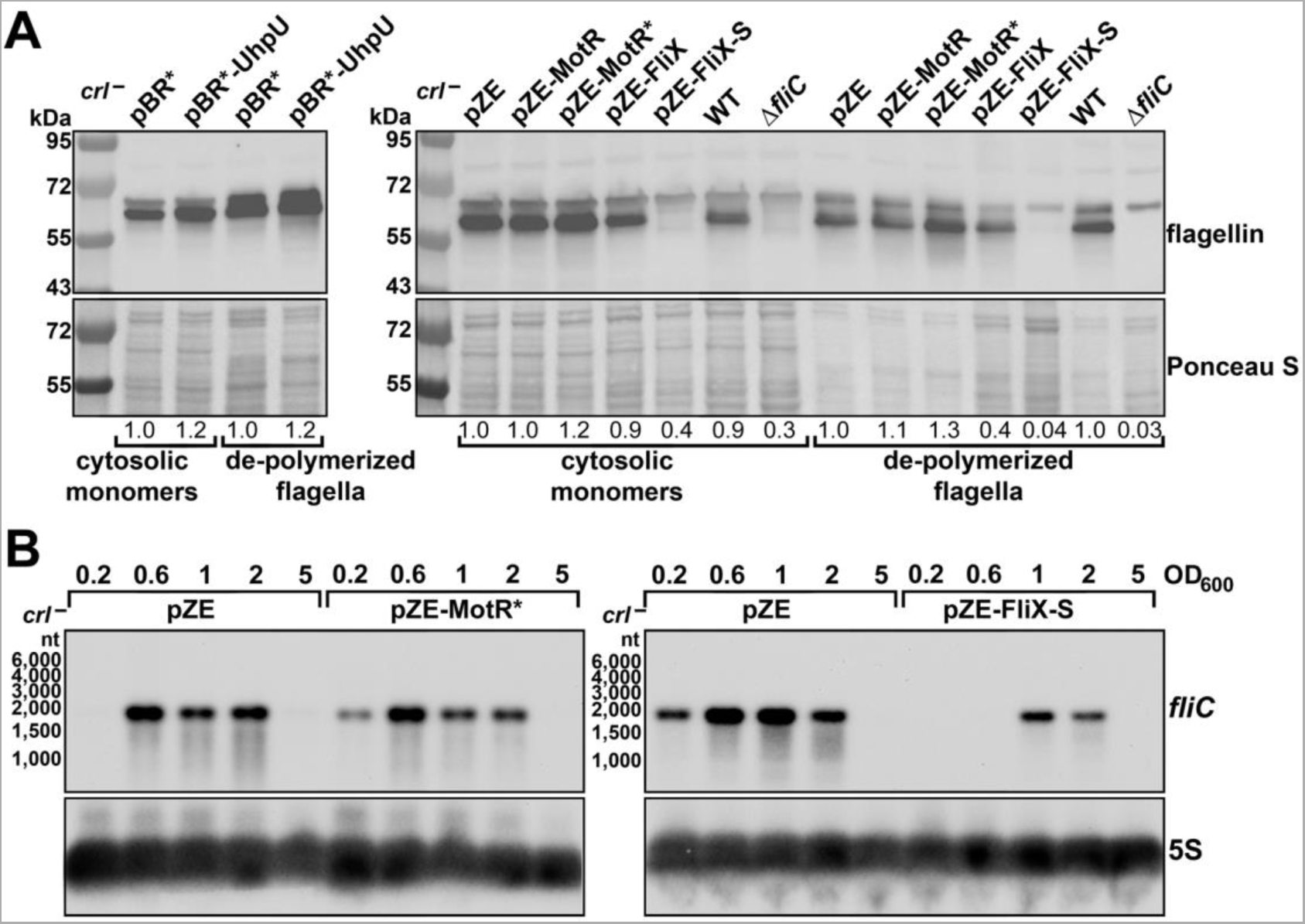
Effects of UhpU, MotR* and FliX-S overexpression on flagellin and *fliC* mRNA levels. (**A**) An expanded view of the immunoblot analysis shown in *Figure 4B*. UhpU and MotR overexpression leads to increased flagellin levels and FliX overexpression leads to reduced flagellin levels. Bacterial cells were fractionated to separate the flagellin cytosolic monomers from the polymerized flagella, and flagellin levels were determined by immunoblot analysis using α-FliC antibody. Numbers represent the fold change relative to the relevant pZE sample. (**B**) Northern blot analysis showing MotR* overexpression increases *fliC* mRNA levels and FliX-S overexpression reduces *fliC* mRNA levels. The 5S RNA served as a loading control. The differences in *fliC* levels in the pZE control samples is due differences in the length of exposure. (C) The membranes used in *Figure 4C* and *Figure 4-figure supplement 1B* were re-probed for the corresponding sRNA overexpressed from plasmid.

**Figure 4—figure supplement 2.**
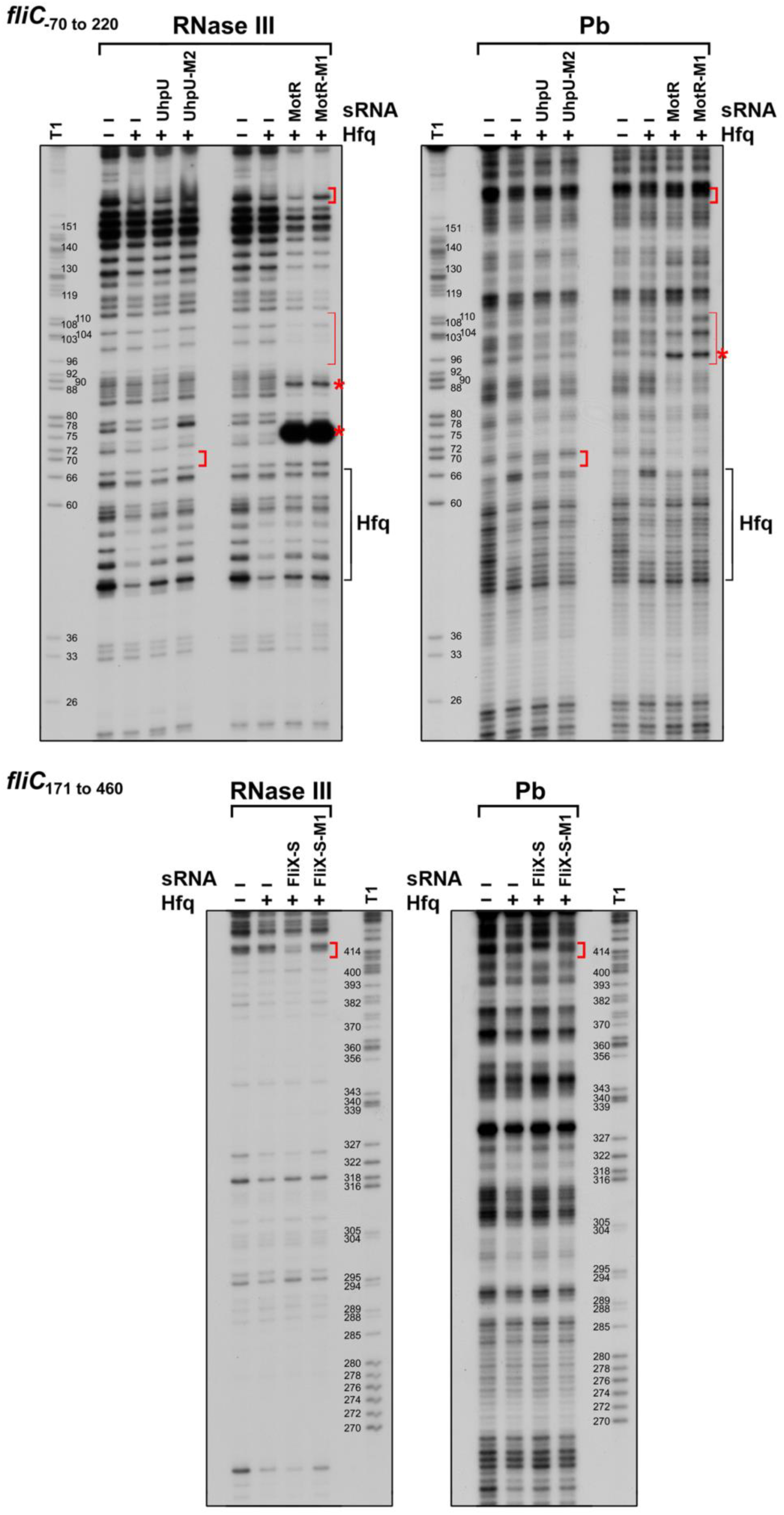
*In vitro* structural probing of interaction between UhpU, MotR and FliX sRNAs with *fliC* mRNA. *In vitro* transcribed and ^32^P-labeled *fliC* fragments (-70 to 220 and 171 to 460 relative to the AUG of the *fliC* open reading frame) were treated with RNase III for 1.5 min or lead for 10 min with or without Hfq and unlabeled UhpU, UhpU-M2, MotR, MotR-M1, FliX-S and FliX-S-M1, and separated, alongside T1 ladder, on a sequencing gel. Changes in cleavage patterns due to the presence of an sRNA, which overlap regions of predicted base pairing, are indicated by the thick red brackets. Asterisks mark increased cleavage, possibly suggesting a change in folding, in the presence of either WT or mutant MotR, and the thin red brackets indicate a possible second site of MotR base pairing. Changes in the cleavage patterns due to Hfq binding are indicated by black brackets. Based on more relative cleavage with RNase III and less cleavage with lead, the *fliC*_-70 to 220_ is more double-stranded than the *fliC*_171-460_ fragment. Numbering is from AUG of *fliC* CDS.

**Figure 5—figure supplement 1.**
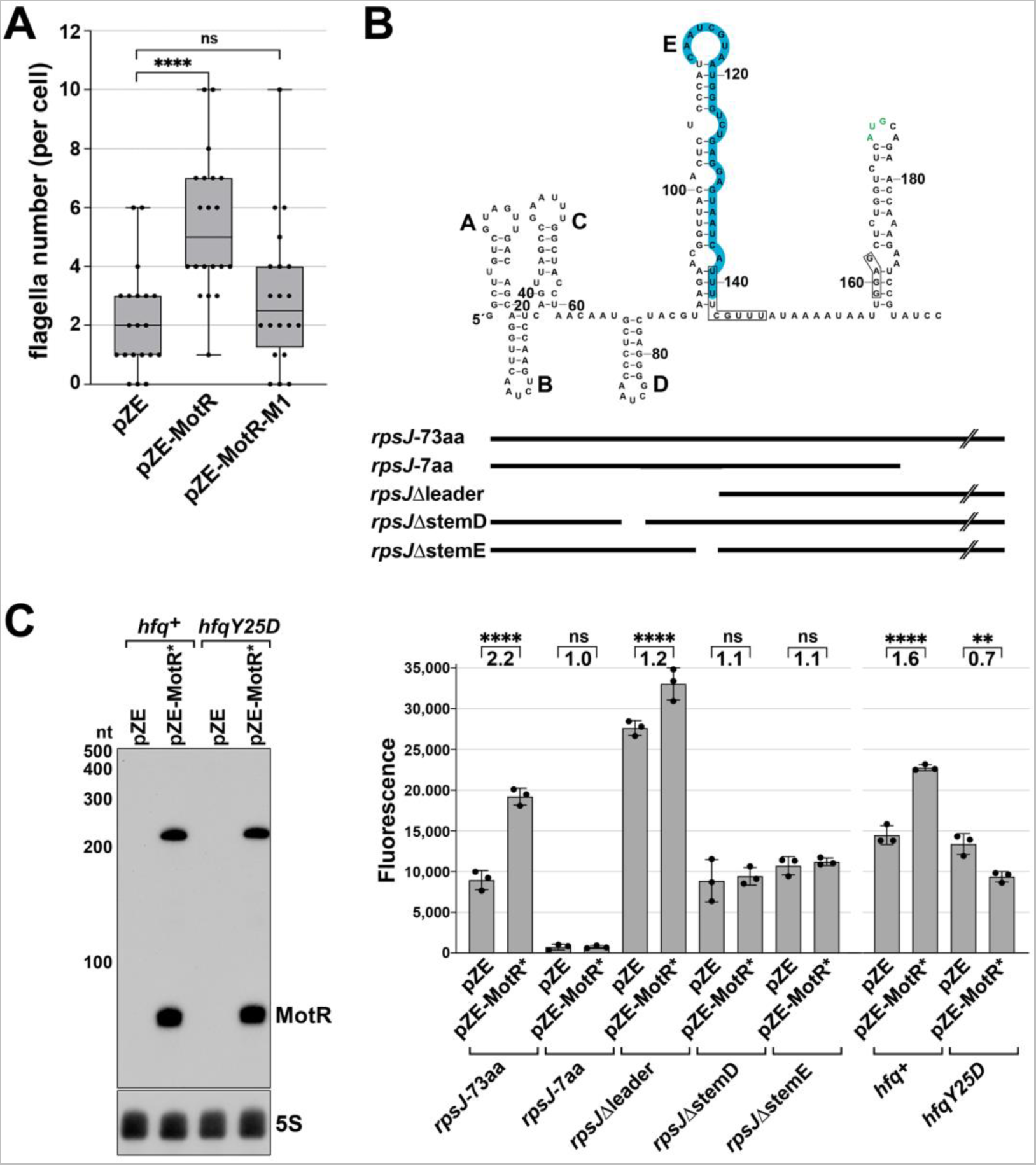
Effects of MotR mutants on flagella number and *rpsJ* expression. (**A**) MotR-M1 overexpression does not increase flagella number. Quantification of the number of flagella per cell by electron microscopy analysis for WT (GSO983) cells harboring the plasmids as indicated in the figure. One-way ANOVA comparison was performed to calculate the significance of the change in flagella number (ns = not significant, **** = P<0.0001). Box plot and error bars descriptions as in Figure 2. (**B**) MotR induces *rpsJ-gfp* levels only when the S10 leader and MotR binding sites are present. S10 leader sequence and secondary structure based on (Zengel *et al*., 2002) (top). Hfq binding region is highlighted in blue. *rpsJ* start codon is labeled in green. The first boxed letters represent S10 leader terminator, and the second boxed letters represent the Shine-Dalgarno sequence. Lines (middle) represent the parts of the sequences that were assayed in the GFP reporter assay in bottom panel. Reporter assays of various *rpsJ-gfp* fusions expressed from pXG10-SF with MotR* expressed from pZE. One-way ANOVA comparison was performed to calculate the significance of the change in GFP signal (ns = not significant, ** = P<0.01, **** = P<0.0001). (**C**) Northern blot analysis of total RNA from *hfq^+^*(GSO614), and *hfqY25D* (GSO1110) grown to OD_600_∼0.2. The *hfq+* and *hfqY25D* strains carried pZE and pZE-MotR*. Similar levels of MotR* RNA were detected for the *hfq*+ and *hfqY25D* mutant backgrounds. Membranes were probed for MotR and then for 5S RNA as a loading control.

**Figure 5—figure supplement 2.**
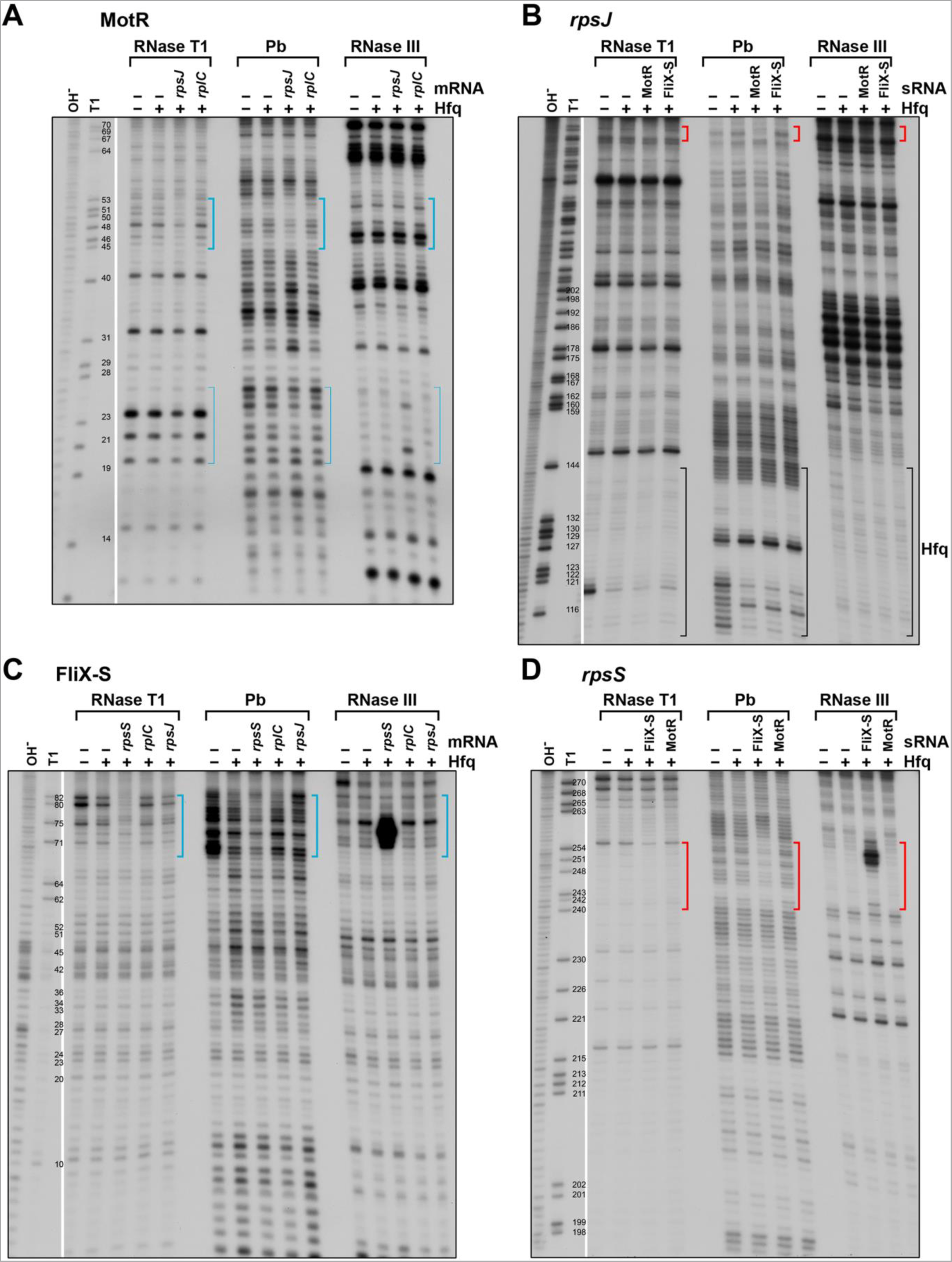
*In vitro* structural probing of interaction between MotR sRNA and *rpsJ* mRNA, and FliX sRNA and *rpsS* mRNA. (**A**) *In vitro* transcribed and ^32^P-labeled MotR was treated with RNase T1 for 10 min, lead for 10 min or RNase III for 1.5 min with or without *rpsJ*, *rplC* and Hfq, and separated, alongside an OH and T1 ladder, on a sequencing gel. Changes in cleavage pattern due to the presence of *rpsJ*, overlapping region of predicted base pairing, are indicated by thick blue brackets. Thin blue brackets indicate a possible second site of base pairing. Numbering is from +1 of MotR. (**B**) *In vitro* transcribed and ^32^P-labeled S10 leader and *rpsJ* was treated with RNase T1 for 10 min, lead for 10 min or RNase III for 1.5 min with or without MotR, FliX-S and Hfq, and separated, alongside an OH and T1 ladder, on a sequencing gel. Changes in cleavage pattern due to the presence of MotR, overlapping region of predicted base pairing, are indicated by thick red brackets, and changes in the cleavage patterns due to Hfq binding are indicated by black brackets. Numbering is from +1 of *rpsJ* mRNA. (**C**) *In vitro* transcribed and ^32^P-labeled FliX-S (with extra 3 nt on its 5’ end as specified in *Supplementary file 3*) was treated with RNase T1 for 10 min, lead for 10 min or RNase III for 1.5 min with or without *rpsS*, *rpsJ*, *rplC* and Hfq, and separated, alongside an OH and T1 ladder, on a sequencing gel. Changes in cleavage pattern due to the presence of *rpsS*, overlapping region of predicted base pairing, are indicated by thick blue brackets. Numbering is from +1 of FliX. (**D**) *In vitro* transcribed and ^32^P-labeled *rpsS* was treated with RNase T1 for 10 min, lead for 10 min or RNase III for 1.5 min with or without MotR, FliX-S and Hfq, and separated, alongside an OH and T1 ladder, on a sequencing gel. Changes in cleavage pattern due to the presence of FliX-S, overlapping region of predicted base pairing, are indicated by thick red brackets. Numbering is from AUG of *rpsS* CDS.

**Figure 5—figure supplement 3.**
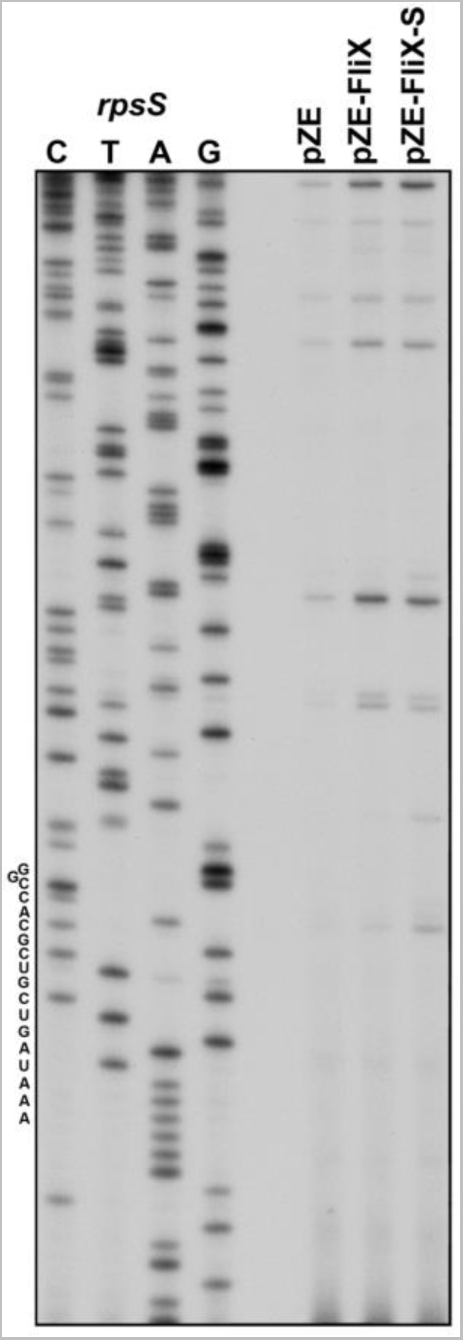
*In vivo* effects of FliX and FliX-S overproduction on *rpsS* mRNA. Total RNA was extracted from a WT strain (GSO983) harboring the indicated plasmids at OD_600_ ∼ 0.6 and then subject to primer extension analysis using a primer hybridizing downstream of FliX binding site on the *rpsS* mRNA.

**Figure 6—figure supplement 1.**
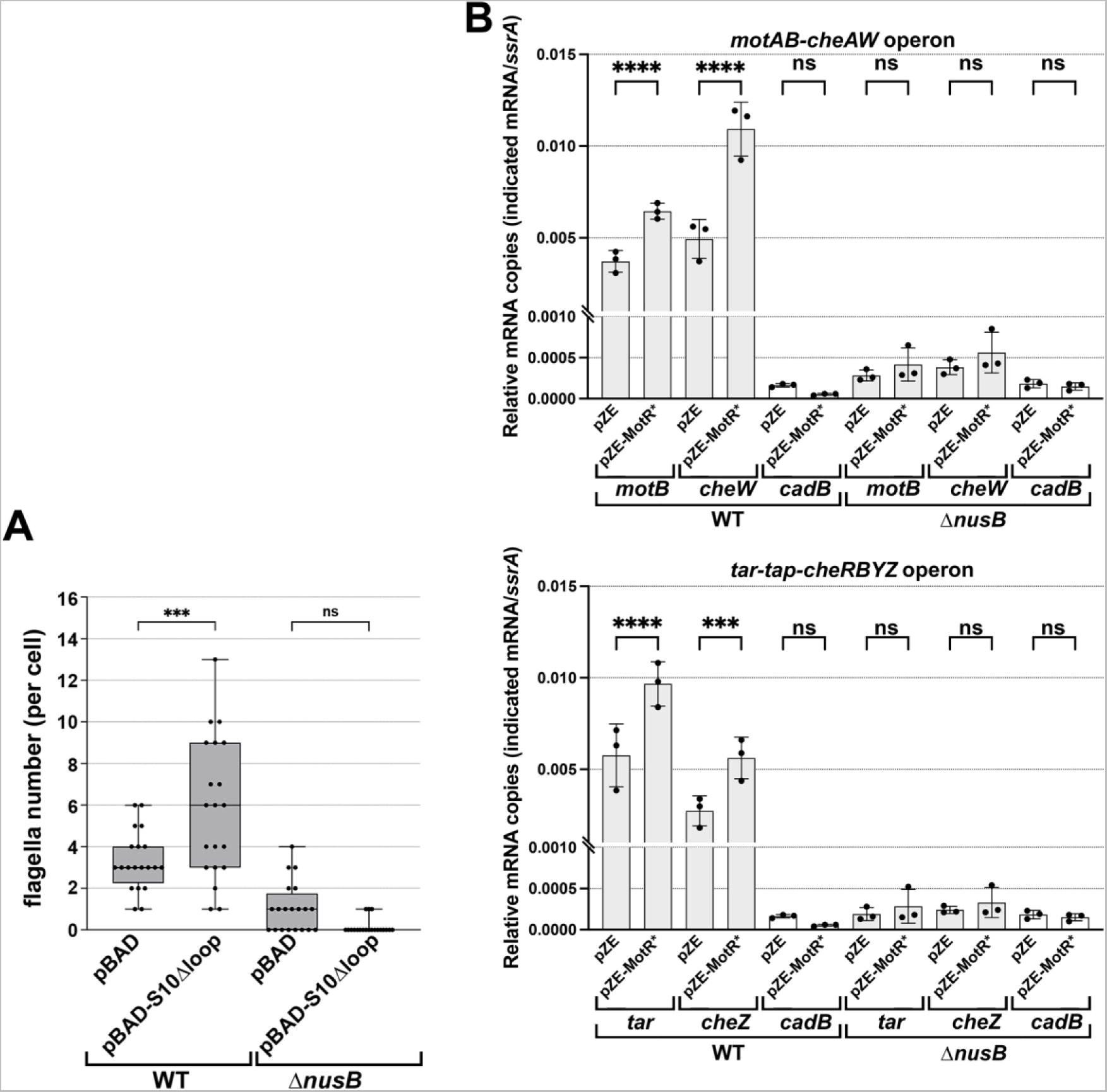
Effects of MotR* and S10Δloop overexpression are lost in Δ*nusB* background. (**A**) S10Δloop effect is eliminated in Δ*nusB* background. The number of flagella per cell detected by electron microscopy were counted for WT (GSO983) or Δ*nusB* (GSO1077) cells harboring the indicated plasmids. Flagella were counted for 20 cells (black dots), and a one-way ANOVA comparison was performed to calculate the significance of the change in flagella number (ns = not significant, *** = P<0.001). Box plot and error bars descriptions as in Figure 2. (**B**) MotR* effect on flagellar operons is eliminated in Δ*nusB* background (GSO1077). MotR* was expressed from pZE plasmid and the levels of *motB*, *cheW*, *tar*, *cheZ*, *ssrA* and *cadB* were monitored in comparison to their levels in the pZE control vector by RT-qPCR. *cadB* served as a non-flagellar gene control and *ssrA* served as a reference gene. Experiments were done in 3 biological replicates and one-way ANOVA comparison was performed to calculate the significance of the change in mRNA levels (ns = not significant, *** = P<0.001, **** = P<0.0001).

**Figure 7—figure supplement 1.**
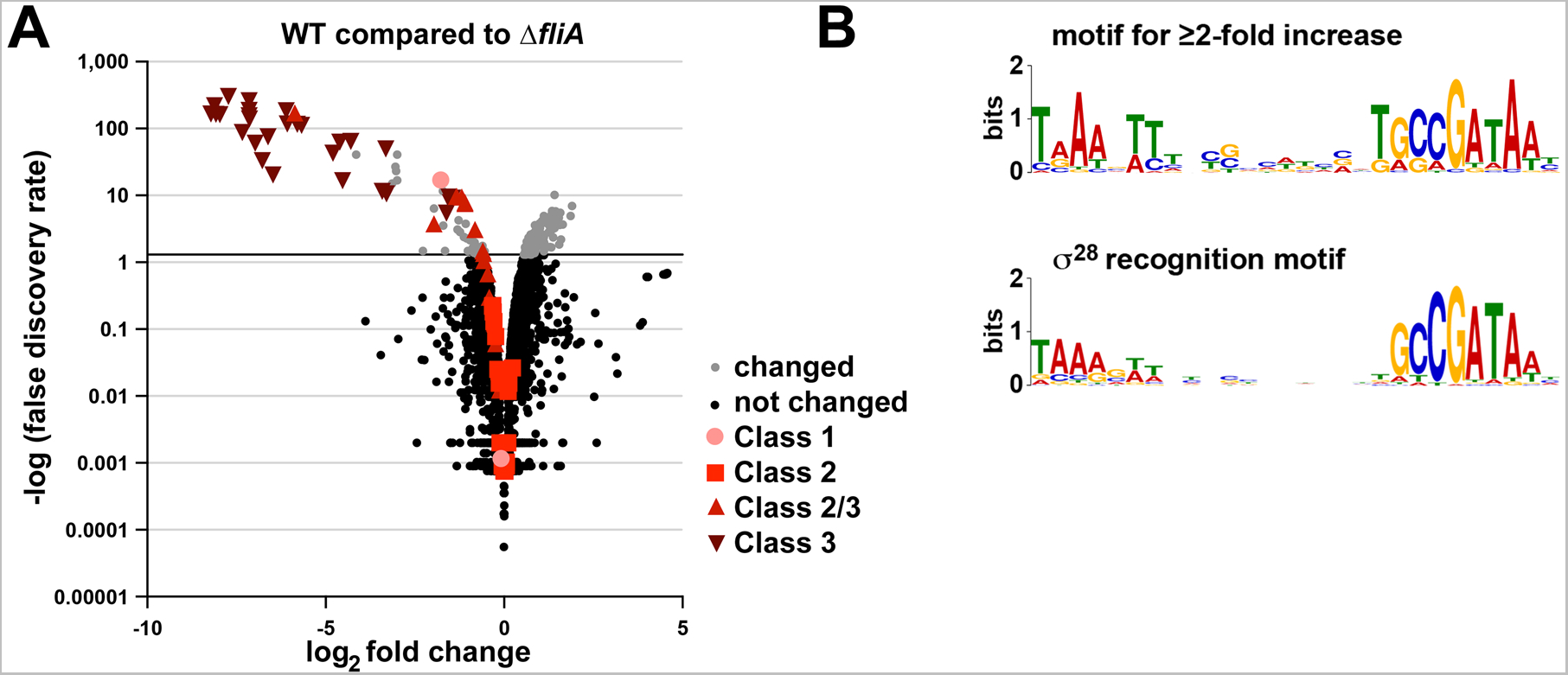
Overlap in MotR* overexpression profile with σ^28^ regulon. (**A**) Deletion of *fliA* reduces σ^28^-dependent genes. RNA-seq data for Δ*fliA* from (Fitzgerald *et al*., 2014) was reanalyzed. Differential expression analysis was conducted with DESeq2 and threshold for differentially expressed transcripts was set to adjusted value of p < 0.05. Red symbols represent flagella regulon genes as indicated on the graph. (**B**) On top is the motif found for promoters of transcription units for genes whose expression increased the most (FDR = 0.05 and ≥2 fold) upon MotR* overexpression (E-value: 5.1e^-14^). Of the 70 total sequences analyzed, 25 contain the motif. The bottom motif corresponds to a σ^28^ recognition motif (Fitzgerald et al. 2014).

**Figure 7—figure supplement 2.**
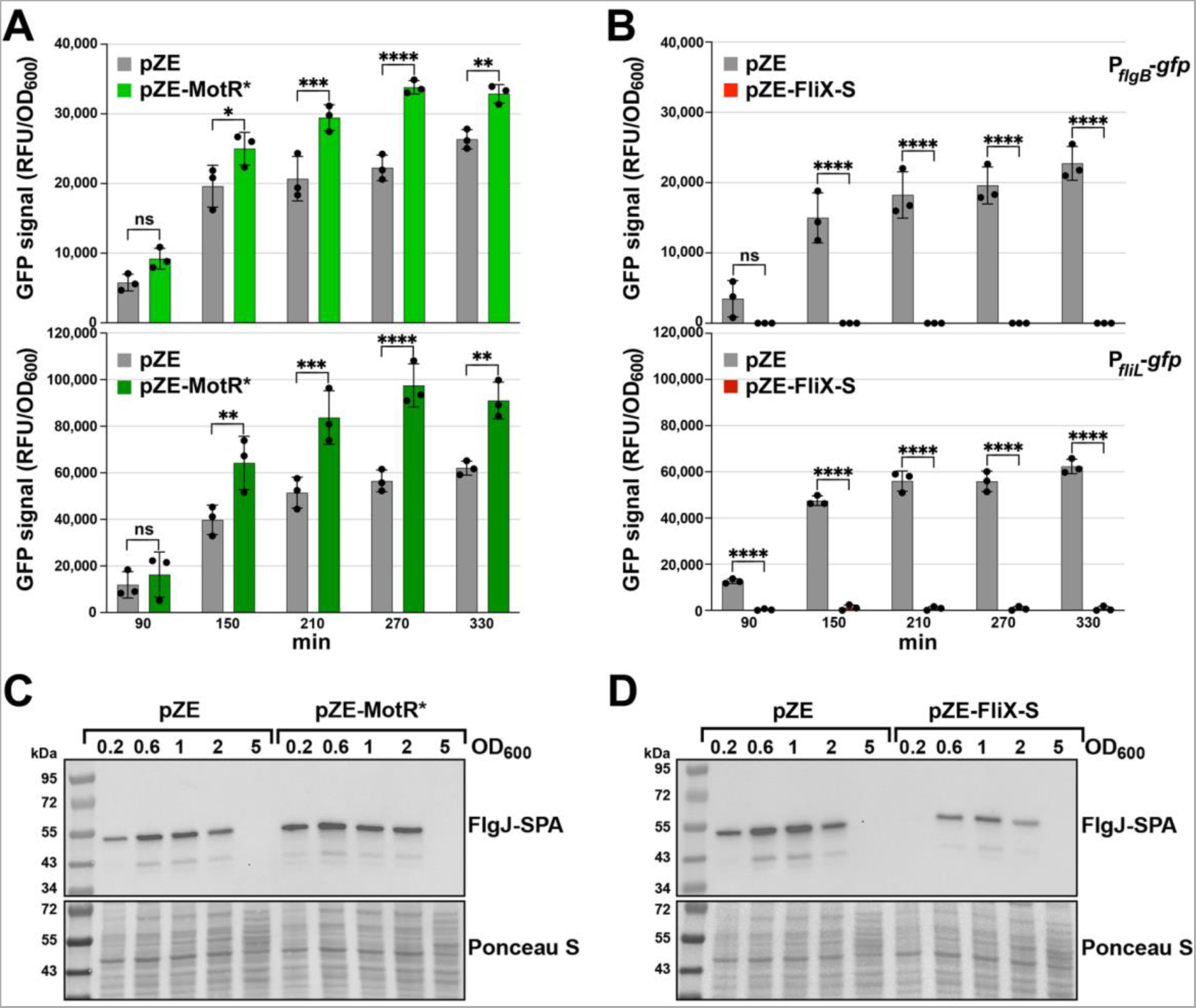
Effects of MotR* and FliX-S overexpression on P*_flgB_*-*gfp,* P*_fliL_*-*gfp* and FlgJ-SPA expression. (**A**) MotR* overexpression increases the activity of GFP fusions to P_flgB_ and P_fliL_. The activity of the promoters was monitored for 330 min by measuring the GFP signal and dividing it with the culture OD_600nm_ in the presence of MotR* expressed from pZE plasmid. (**B**) FliX-S overexpression decreases the activity of GFP fusions to P_flgB_ and P_fliL_. The activity of the promoters was monitored for 330 min by measuring the GFP signal and dividing it with the culture OD_600nm_ in the presence of FliX-S expressed from pZE plasmid. (**C**) MotR* increases the levels of SPA-tagged FlgJ. FlgJ-SPA levels across growth of WT (GSO1080) cells in the presence of MotR* expressed from pZE plasmid were determined by immunoblot analysis using α-FLAG antibody. (**D**) FliX-S reduces the levels of SPA tagged-FlgJ. FlgJ-SPA levels across growth of WT (GSO1080) cells in the presence of FliX-S expressed from pZE plasmid were determined by immunoblot analysis using α-FLAG antibody. For (**A**) and (**B**), three biological repeats are shown in the graph. One-way ANOVA comparison was performed to calculate the significance of the change in GFP signal (ns = not significant, * = P<0.05, ** = P<0.01, *** = P<0.001, **** = P<0.0001). The experiments presented in 7C and *Figure 7— figure supplement 2B*, and in 7D and *Figure 7—figure supplement 2A*, were carried out on same day, respectively, and the same pZE samples are shown. For (**C**) and (**D**), the Ponceau S-stained membrane serves as a loading control.

**Figure 8—figure supplement 1.**
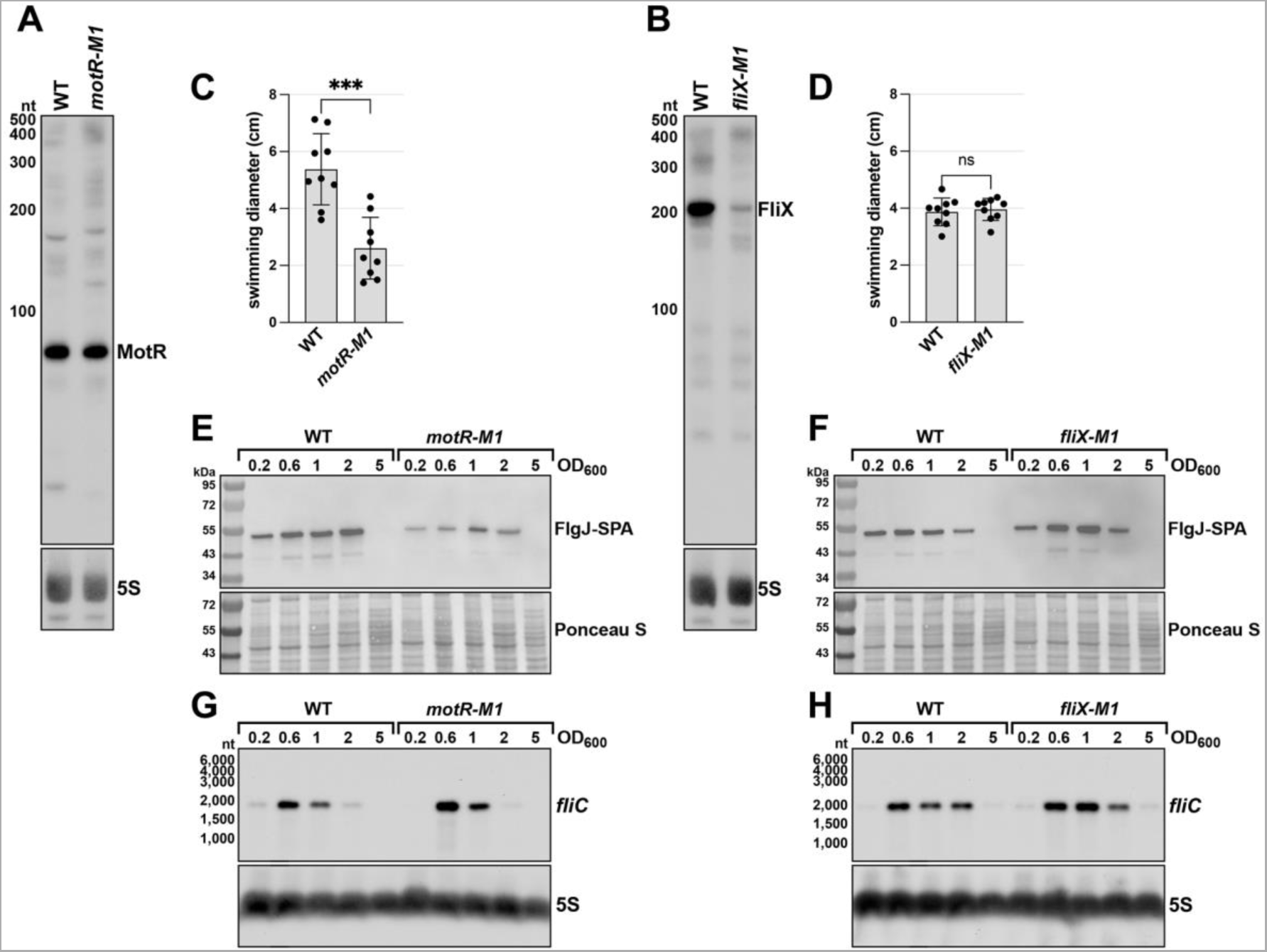
Effects of chromosomal *motR-M1* and *fliX-M1* mutations. (**A**) Northern blot analysis of total RNA from WT (GSO1088), and *motR-M1* (GSO1087) grown to OD_600_∼1.0. The levels of MotR-M1 expressed from the chromosome are comparable to those of WT MotR. Membranes were probed for MotR and then for 5S RNA as a loading control. (**B**) Northern blot analysis of total RNA from WT (GSO983), and *fliX-M1* (GSO1076) grown to OD_600_∼1.0. The levels of full length FliX-M1 expressed from the chromosome are lower than WT FliX. Membranes were probed for FliX and then for 5S RNA as a loading control. (**C**) Reduced motility in *motR-M1* mutant (GSO1087) compared to corresponding WT strain (GSO1088) based on motility assays with 0.3% agar. (**D**) No change in motility of *fliX-M1* mutant (GSO1076) compared to corresponding WT strain (GSO983) based on motility assays with 0.3% agar. (**E**) Reduced FlgJ-SPA levels in a *motR-M1* mutant. SPA-tagged FlgJ levels across growth in WT (GSO1081) and in *motR-M1* mutant (GSO1082) were monitored. (**F**) Slightly increased in FlgJ-SPA levels in a *fliX-M1* mutant. SPA-tagged FlgJ levels across growth in WT (GSO1080) and in *fliX-M1* mutant (GSO1083) were monitored. Northern blot analysis showing (**G**) delayed expression of *fliC* mRNA in *motR-M1* background and (**H**) advanced expression in *fliX-M1* background.(**H**). Graphs in (**C**) and (**D**) show the average of nine biological replicates, and error bars represent one SD. One-way ANOVA comparison was performed to calculate the significance of the change in motility (ns = not significant, *** = P<0.0001). In (**E**) and (**F**), FlgJ-SPA levels were determined by immunoblot analysis using α-FLAG antibody. The Ponceau S-stained membrane serves as a loading control. In (**G**) and (**H**), *fliC* mRNA levels were determined by northern analysis as in Figure 4. RNA was separated on an acrylamide gel and probed for *fliC* and then for the 5S RNA as a loading control.

**Figure 8—figure supplement 2.**
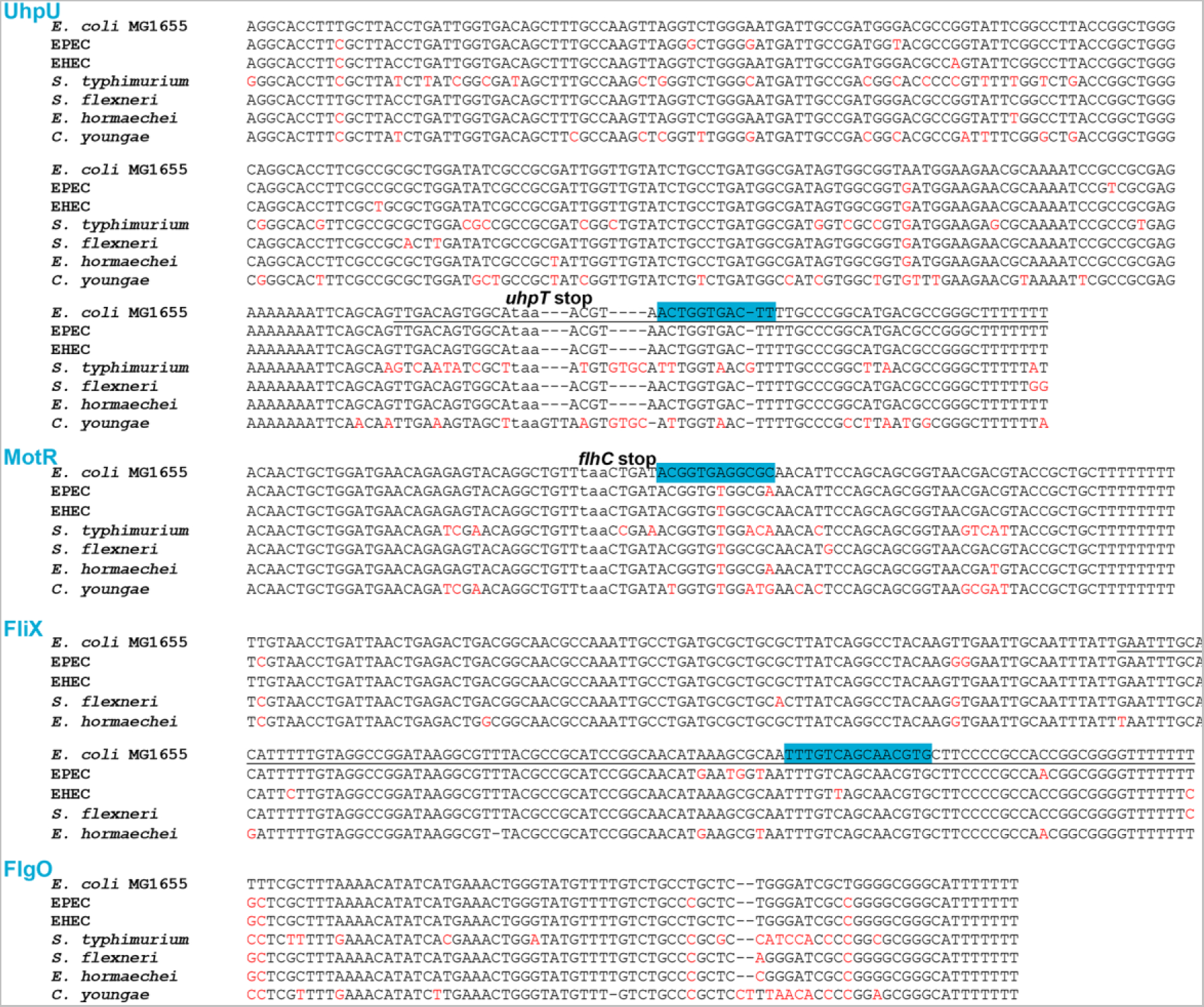
Conservation of σ^28^-dependent sRNAs. σ^28^-dependent sRNAs sequences were tested for their conservation across the Gammaproteobacteria, revealing that their conservation in closely-related Enterobacteriaceael organisms. Sequences from representative bacterial species are aligned as follows. Gammaproteobacteria genomes were downloaded from PATRIC database (Davis *et al*., 2020) and searched for the query sRNAs using BLAST (Altschul *et al*., 1990), filtering for matches with at least 80% identity over 80% of the query sRNA sequence. Genomes containing each sRNA were marked as so on a text file. The process was written as a Nextflow pipeline (Di Tommaso *et al*., 2017) and is available on github: https://github.com/asafpr/sRNA_finder. The analysis results were confirmed by aligning the sRNAs sequences to representative Gamma proteobacteria species. The *uhpT* and *flhC* stop codons are indicated in lower case. Regions highlighted in blue reflect the suggested seed sequence of *uhpU* (Melamed *et al*., 2016) and the regions that base pair with the targets tested in this study for *motR* and *fliX*. The *uhpU-S* and *fliX-S* sequences are underlined. Nucleotides that differ from the *E. coli* MG1655 sequence are indicated in red. The abbreviations in the figure represent the following species: *Escherichia coli* MG1655 (*E. coli* MG1655), *Escherichia coli* O127:H6 (EPEC), *Escherichia coli* O157:H7 (EHEC), *Salmonella enterica serovar typhimurium* (*S. typhimurium*), *Shigella flexneri* (*S. flexneri*), *Enterobacter hormaechei* (*E. hormaechei*), and *Citrobacter youngae* (*C. youngae*). *S. typhimurium* and *C. youngae* are missing from the alignment for FliX as the *fliX* region was missing in these species.

